# mRNA targeting eliminates the need for the signal recognition particle during membrane protein insertion in bacteria

**DOI:** 10.1101/2022.11.21.517368

**Authors:** Pinku Sarmah, Wenkang Shang, Andrea Origi, Mariya Licheva, Claudine Kraft, Maximilian Ulbrich, Elisabeth Lichtenberg, Annegret Wilde, Hans-Georg Koch

## Abstract

Signal-sequence dependent protein targeting is essential for the spatiotemporal organization of eukaryotic and prokaryotic cells and facilitated by dedicated protein targeting factors, such as the signal recognition particle (SRP). However, targeting signals are not exclusively contained within proteins, but can also be present within mRNAs. By *in vivo* and *in vitro* assays, we show that mRNA targeting is controlled by the nucleotide content and by secondary structures within mRNAs. mRNA binding to bacterial membranes occurs independently of soluble targeting factors, but is dependent on the SecYEG-translocon and YidC. Importantly, membrane insertion of proteins translated from membrane-bound mRNAs occurs independently of the SRP pathway, while the latter is strictly required for proteins translated from cytosolic mRNAs. In summary, our data indicate that mRNA targeting acts in parallel to the canonical SRP-dependent protein targeting and serves as an alternative strategy for safeguarding membrane protein insertion when the SRP pathway is compromised.

## Introduction

Protein transport across membranes is an essential process and depends on largely conserved mechanisms that include dedicated protein translocases and insertases as well as specific protein targeting factors ^1–4^. The Sec translocon is the best characterized protein translocase ^5, 6^ and largely conserved between pro- and eukaryotes ^6, 7^. A high conservation is also seen in the Oxa1 superfamily of protein insertases, which insert membrane proteins into the bacterial cytoplasmic membrane and into organellar membranes in eukaryotes ^8–10^.

In bacteria, most membrane proteins are targeted co-translationally by the conserved signal recognition particle (SRP) and its receptor FtsY to either the Sec translocon or to the Oxa1 superfamily member YidC ^11–15^. SRP binds to the exit of the ribosomal peptide tunnel and scans the tunnel for membrane protein substrates ^16–20^. Once bound to the emerging signal anchor sequence of the nascent membrane protein, SRP targets the ribosome-associated nascent chain (RNC) to FtsY ^21, 22^, which is bound to the cytoplasmic loops of SecY and YidC^23–30, 31, 32^. Upon SRP-FtsY interaction, the RNC docks onto SecY or YidC ^20, 33^ and ongoing translation together with the thermodynamically favoured lipid partitioning of transmembrane domains facilitates membrane insertion ^34, 35^.

While signal sequence based protein targeting is well established ^36–41^, the contribution of mRNA targeting to protein transport is largely unknown. Imaging techniques have revealed distinct mRNA localization patterns and the membrane enrichment of mRNAs encoding for membrane proteins in bacteria, such as *Escherichia coli* or *Lactococcus lactis* ^42–45^. However, whether mRNA targeting depends exclusively on sequence and structural information within the mRNA or relies also on information within the encoded protein is unclear ^43, 45, 46^. Translation-independent membrane targeting of mRNA requires an RNA zip code that determines its cellular localization. The uracil content within the mRNA potentially provides such a zip code, because bioinformatics revealed a uracil bias in transcripts encoding for membrane proteins ^47, 48^. This is supported by data showing that increasing the uracil content in a cytosolic transcript enhanced membrane localization ^43^, but the corresponding mRNA receptors are still unknown. Cold-shock proteins have been suggested to be involved in mRNA targeting ^49, 50^ and a possible role of FtsY was also proposed ^51^, but in general, the roles of SRP and FtsY in handling membrane proteins that are translated from membrane-bound mRNAs are unknown.

In the current study, we combined *in vivo* imaging with biochemical assays for further exploring the contribution of mRNA targeting to membrane protein insertion in bacteria. *In vivo* localization studies demonstrate that in addition to the uracil content the secondary structure of an mRNA also contributes to membrane binding. *In vitro* mRNA binding assays revealed that sequence-specific membrane binding is determined by the SecYEG and YidC content. Intriguingly, *in vitro* assays demonstrate that protein insertion into the membrane occurs independently of the SRP pathway, when the protein is translated from already membrane-bound mRNAs. In contrast, when produced from cytosolic mRNAs, insertion strictly dependent on SRP and FtsY for insertion. Our data indicate that translation-independent mRNA targeting provides an SRP-independent mechanism for membrane protein insertion in bacteria.

## Results

### Determinants of mRNA membrane localization

In most *in vivo* studies, mRNA localization is visualized by using the MS2 reporter system, which employs the fluorescently labelled phage protein MS2. MS2 binds to a hexa-repeat stem-loop sequence that is fused to the 3’-end of a target mRNA ^42, 43, 48, 51–53^.

In the current study, the MS2 reporter system was employed to analyse the localization of mRNAs encoding for the single-spanning membrane protein YohP or the multi-spanning membrane protein SecY ^53^. For determining mRNA localization independently of translation, the Shine-Dalgarno sequence of the analysed mRNAs was removed. In the absence of any mRNA, MS2-Venus preferentially localized to the bacterial cytoplasm and showed some clustering, which likely reflects the formation of aggregates or inclusion bodies (Fig. 1A). Inclusion bodies contain a significant amount of lipids ^54^, which probably explains why these clusters were also visible when cells were stained with the membrane-specific dye Nile red. In contrast, when MS2-Venus was co-expressed with *yohP* mRNA fused to the hexa-repeat stem loop, MS2-Venus was almost exclusively membrane localized (Fig. 1A). The *secY* mRNA, which encodes for the 48 kDa SecY subunit of the SecYEG translocon ^55^, showed also membrane enrichment, indicating that membrane localisation is not a particular feature of small mRNAs. In contrast, the mRNA of the cytosolic protein BglB primarily localized to the cytoplasm (Fig. 1A). mRNA localization was quantified in multiple cells (n >60-80 cells) using the Jenson-Shannon divergence (JSD), which is a measure for the divergence of two distributions ^56^ (Fig. 1B & Fig. S1A). A low JSD value indicates that Nile red and the mVenus signals show a similar distribution within the cell, *i.e.* a predominant membrane localization of MS2-Venus. In contrast, a high JSD value reveals a cytosolic localization of MS2-Venus. The lowest JSD value was observed for mVenus/Nile red in the presence of the *yohP* mRNA, followed by the *secY* mRNA (Fig. 1B). The JSD in the presence of the *bglB* mRNA or in the absence of any target mRNA were considerably higher, indicating a low overlap between the Nile red and MS2-venus distributions.

**Figure 1:**
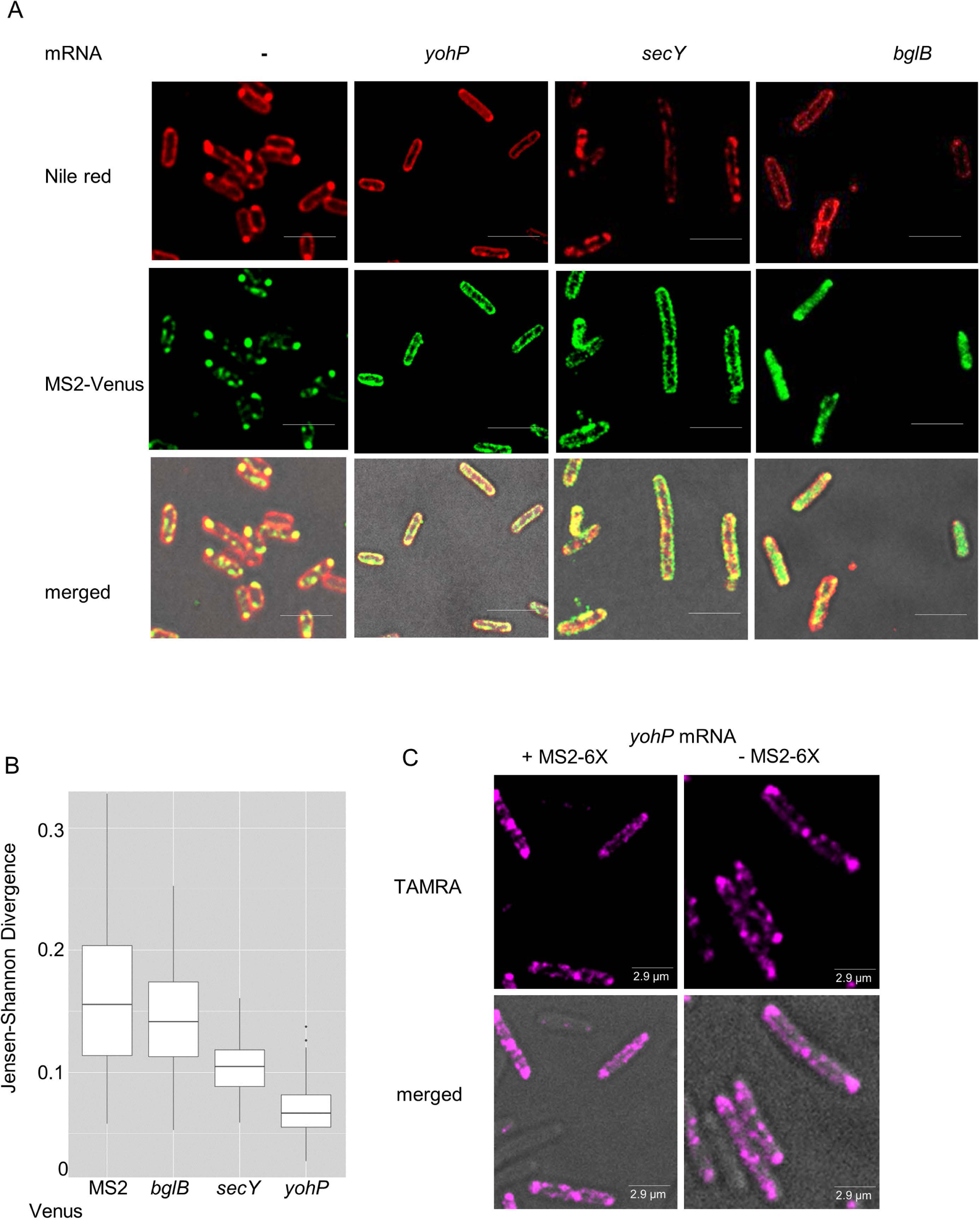
Translation-independent membrane enrichment of mRNAs encoding for membrane proteins. **(A)** *E. coli* wild type cells expressing just pBad24-MS2-Venus or together with a plasmid encoding either *yohP*, *secY* or *bglB*, each with a deleted ribosome binding site and a hexa-repeat MS2 stem loop sequence at the 3’-UTR. mRNA expression was induced at OD_600_ 0.5 with 1 mM IPTG and MS2-Venus was induced 1h after IPTG addition with 0.2 mM arabinose. Of each culture, 10 µL solution were placed on a sterile glass bottom dish. Imaging was performed with the Delta Vision^TM^ Ultra microscope (exposure 0.4 sec for mVenus at 32% laser intensity and 0.075 sec for bright field at 5% laser intensity) and 3µm Z-scans were recorded with an interval of 1µm. The images were taken at the focal point. The image was developed with the *ImageJ Fiji* software. Nile red was used as membrane stain and staining was monitored with an exposure time of 0.1 sec at a laser intensity of 10%. The scale bar refers to 2.5 µM. **(B)** Jensen-Shannon divergence (JSD) plot for quantifying the correlation between Nile red distribution and mVenus distribution (See also Fig. S1). The JSD was evaluated with scores between 0 (identical distribution) and 1 (maximally different distribution) and is based on scoring 60-80 individual cells. (**C**) RNA-FISH of *E. coli* cells expressing *yohP* mRNA with or without the MS2 stem-loop sequence. A set of 19 oligonucleotides against the *yohP* sequence were linked to the fluorescent probe TAMRA (absorption maximum 522, emission maximum 576 nm). Imaging was performed as in (A) with 0.1 sec exposure time and 100% laser intensity. (See also Figure S1 and Figure S6).

The *secY* and *yohP* constructs used for *in vivo* localization studies only contained the respective coding sequence, but not the respective 5‵- or 3‵- untranslated regions (UTRs), indicating that the targeting information is retained within the coding sequence. However, the *yohP* coding sequence is rather short (81 nucleotides) and the absence of 5‵- and 3‵-UTRs could influence membrane localization. This was tested by analysing a *yohP* mRNA that contained both UTRs, but this did not change its membrane localization (Fig. S1B & S1C). Whether the attachment of the MS2 stem loop to the *yohP* mRNA influences membrane localization was analysed by RNA *fluorescence in situ hybridization* (FISH) experiments with TAMRA (5-carboxytetramethylrhodamine)-labelled *yohP* probes (Figs. 1C & Fig. S1D). Cells expressing the *yohP* mRNA containing the 5′ and 3′-UTRs with or without the MS2 stem-loop showed an identical localization pattern with most of the fluorescent signal at the membrane (Fig. 1C), indicating that membrane localization of the *yohP* mRNA is not significantly influenced by the MS2 stem loop.

The *yohP* mRNA lacking the 5‵- and 3‵-UTRs, but containing the stem loops was then chosen as model mRNA for identifying possible targeting signals within its coding sequence. Several *yohP* mRNA variants were constructed, which all maintained the reading frame (Fig. S2), but differed in nucleotide composition, secondary structure or mRNA length.

When the uracil content of the *yohP* mRNA was increased to more than 50% of the total nucleotide content, enhanced membrane binding was observed (Fig. 2A; Fig. S3A). In contrast, a two-fold increase of the cytosine content reduced membrane localization and induced clustering of MS2-venus. A similar effect was observed when the adenine content was increased. Finally, increasing the guanine content also enhanced membrane binding of the *yohP* mRNA, but also showed some clustering (Fig. 2A; Fig. S3A).

**Figure 2:**
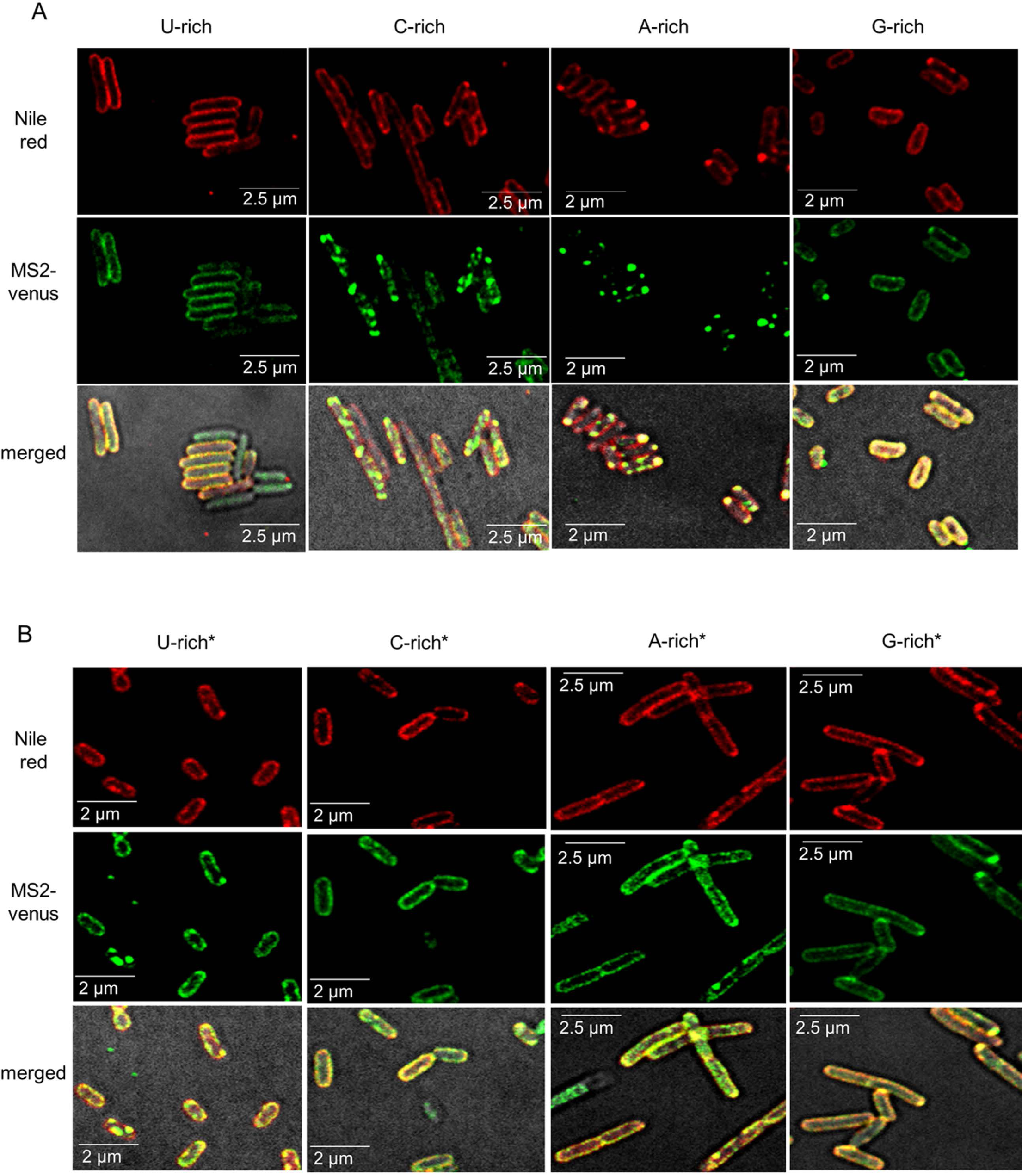
Membrane localization of the *yohP* mRNA is dependent on both nucleotide sequence and secondary structure. **(A)** The wild type *yohP* sequence was modified by increasing either the uracil content (U-rich), the cytosine content (C-rich), the guanine content (G-rich) or the adenine content (A-rich). Imaging was performed as described in the legend of Fig. 1. **(B)** As in (A), but the mRNA variants with increased U-, C-, A- or G-content were designed with only minor influence on the predicted secondary structure of the *yohP* mRNA (U-rich*, C-rich*, G-rich*, A-rich*). The exact nucleotide sequences, compositions and secondary structures are displayed in Fig.S2 and a JSD plot in Fig. S3A.

*In silico* analyses predict the presence of two stem loops in the *yohP* mRNA (nt 10-25 and 30-78), which are altered in the U-, C-, A-, G-rich *yohP* variants (Fig. S2C). It is therefore difficult to determine whether nucleotide composition, secondary structures, or both determine membrane localization. This was investigated by analysing membrane binding of *yohP* variants that largely maintained the overall predicted mRNA structure, despite variations in the nucleotide content (Fig. S2C). Maintaining the secondary structure in uracil-enriched or guanine-enriched *yohP* variants (U-rich*, G-rich*) had no major influence on membrane localization, as compared to the original U- and G-rich variants. In contrast, maintaining the secondary structure for the cytosine- or adenine-enriched *yohP* mRNAs (C-rich*, A-rich*) enhanced their membrane localization, in comparison to the original C- and A-rich variants (Figs. 2B, S3A).

The possibility that differences in transcript abundance influence membrane localization of the *yohP* mRNA was addressed by Northern blot experiments. There were some variations in the *in vivo* abundance of the different mRNAs (Fig. S3) and in particular a higher abundance of the G-rich *yohP* mRNA, but besides this we did not detect drastic differences, which could explain the different localization pattern.

The contribution of secondary structures to membrane localization of the *yohP* mRNA was further studied by generating deletion variants (Figs. 3A, S2). Deleting either nucleotides 4-30 or 10-27 prevented membrane localization and induced MS2 clustering (Figs. 3B; S2; S3F). YohP mRNAs lacking nucleotides 46-60 maintained their ability to interact with the membrane, but induced cell elongation (Fig. 3B). These deletions also reduced the mRNA length and therefore additional variants were tested in which the loops were replaced with random sequences. Replacing the deleted nucleotides 4-30 with a random sequence (Fig. S2) did not interfere with membrane targeting, as compared to the wild type *yohP* mRNA (Fig. 3C), and a similar observation was made when nucleotides 10-27 were replaced. However, these replacements also increased cell length, which was also observed when nucleotides 46-60 were replaced. In this latter mRNA, membrane localization was reduced as well.

**Figure 3:**
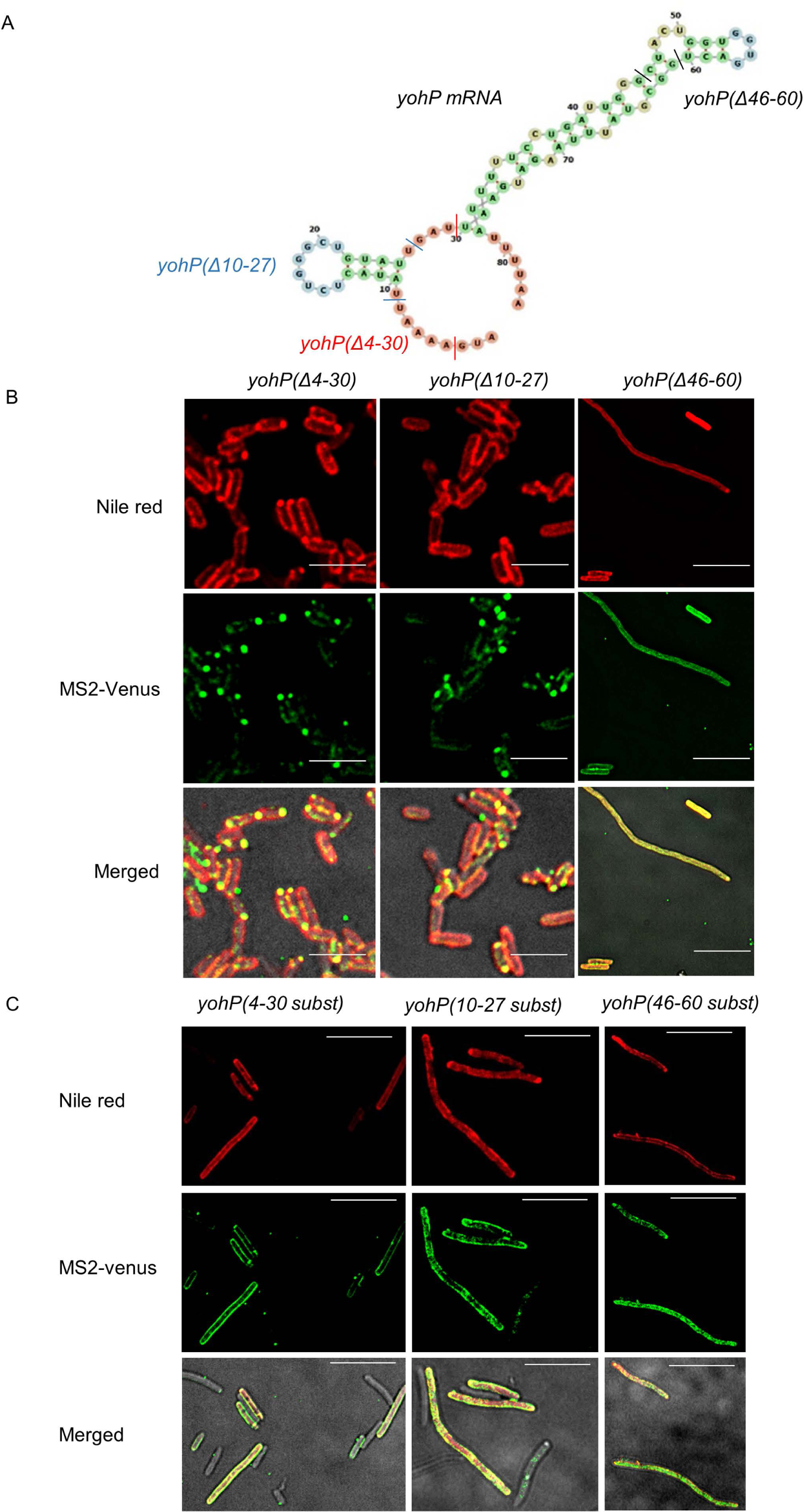
Deletion or replacements of the predicted loops of the *yohP* mRNA impair membrane binding. **(A)** Secondary structure prediction of the *yohP* mRNA using the *RNAfold* webserver (http://rna.tbi.univie.ac.at/cgi-bin/RNAWebSuite/RNAfold.cgi). The predicted loops and their deletions are indicated by colored lines. **(B)** Imaging of *E. coli* cells expressing pBad-24-MS2-Venus together with the indicated stem-loop deletions, as described in the legend of Fig. 1. **(C)** As is (B), but the sequence of the predicted stem loops were replaced with random sequences. All sequence information is shown in Fig. S2 and the quantification of the data in (B) in Fig. S3C. The scale bar refers to 2.5 µM.

The observation that the uracil content is important for membrane localization of bacterial mRNAs is in line with previous proposals ^43, 47, 49^. In eukaryotes, it has been shown that increasing either the uracil or cytosine content enhanced mRNA binding to the ER membrane ^57^, but comparing membrane localization of the C-rich and C-rich* variants indicates that enhanced membrane binding of cytosine-enriched mRNAs is only observed when the overall secondary structure of the mRNA is maintained. The length of the mRNA is apparently also important, because replacing nucleotides 10-27 with a random sequence restored membrane localization. However, this could be primarily important for small mRNAs, such as the *yohP* mRNA. Finally, although all tested mRNAs lacked the Shine-Dalgarno sequence and were not translated *in vitro* as exemplified for the wild type *yohP* (Fig. S4A), some *yohP* mRNA variants influenced cell length/volume, but the underlying mechanism was not further explored in the current study.

### The SecYEG translocon and YidC act as potential mRNA receptors at the bacterial membrane

RNAs can bind to phospholipid surfaces ^58–61^ and the lipid surface of the membrane could potentially be sufficient for mRNA binding. This was analysed *in vitro* by incubating *in vitro* transcribed ^32^P-labeled *yohP* mRNA with liposomes of different lipid composition, followed by ultracentrifugation for separating bound mRNA from non-bound mRNA (Fig. S4B). *In vitro* transcription of *yohP* resulted in two distinct bands (Fig. S4B), probably because the six consecutive MS2 stem-loops remain partially folded in the denaturing buffer due to their high stability ^52, 62^. In the absence of liposomes, the mRNA was found exclusively in the supernatant (S) after centrifugation (Fig. S4B), while in the presence of liposomes mirroring the natural *E. coli* membrane lipid composition (70% PE, 25% PG, 5% CL), a large portion of the mRNA was found in the pellet fraction (Fig. S4B). Variations in the lipid composition (Fig. S4B) had only a small impact on mRNA binding, with the exception of liposomes with 100% phosphatidyl glycerol (PG), which showed less mRNA binding (Fig. S4C). A previous study had shown RNA-induced liposome aggregation, which was not observed for PG-containing liposomes ^58^. Thus, although our data support the interaction of mRNA with phospholipid surfaces, further interpretation is complicated by mRNA-induced liposome aggregation, which influences the sedimentation assay. In eukaryotes, mRNA binding proteins bind certain mRNAs prior to translation and protein transport ^63–65^. Whether such proteins are also present in the bacterial membrane was analysed by repeating the mRNA binding assay in the presence of sucrose-gradient purified inner membrane vesicles of *E. coli* (INVs). In the absence of INVs, the *yohP* mRNA was found in the supernatant while in the presence of wild type INVs, a portion of the mRNA was recovered from the pellet fraction (Fig. 4A, B). The incubation of the *yohP* mRNA with INV resulted in an additional band (Fig. 4A, *) and generally in more diffuse bands. The diffuse bands are likely an indication for mRNA degradation by the membrane-attached *E. coli* RNA degradosome ^66^, which could not be completely prevented even at higher concentrations of the RNase inhibitor RNasin. The additional band potentially reflects incompletely denatured *yohP* mRNA that did not fully enter the gel.

**Figure 4:**
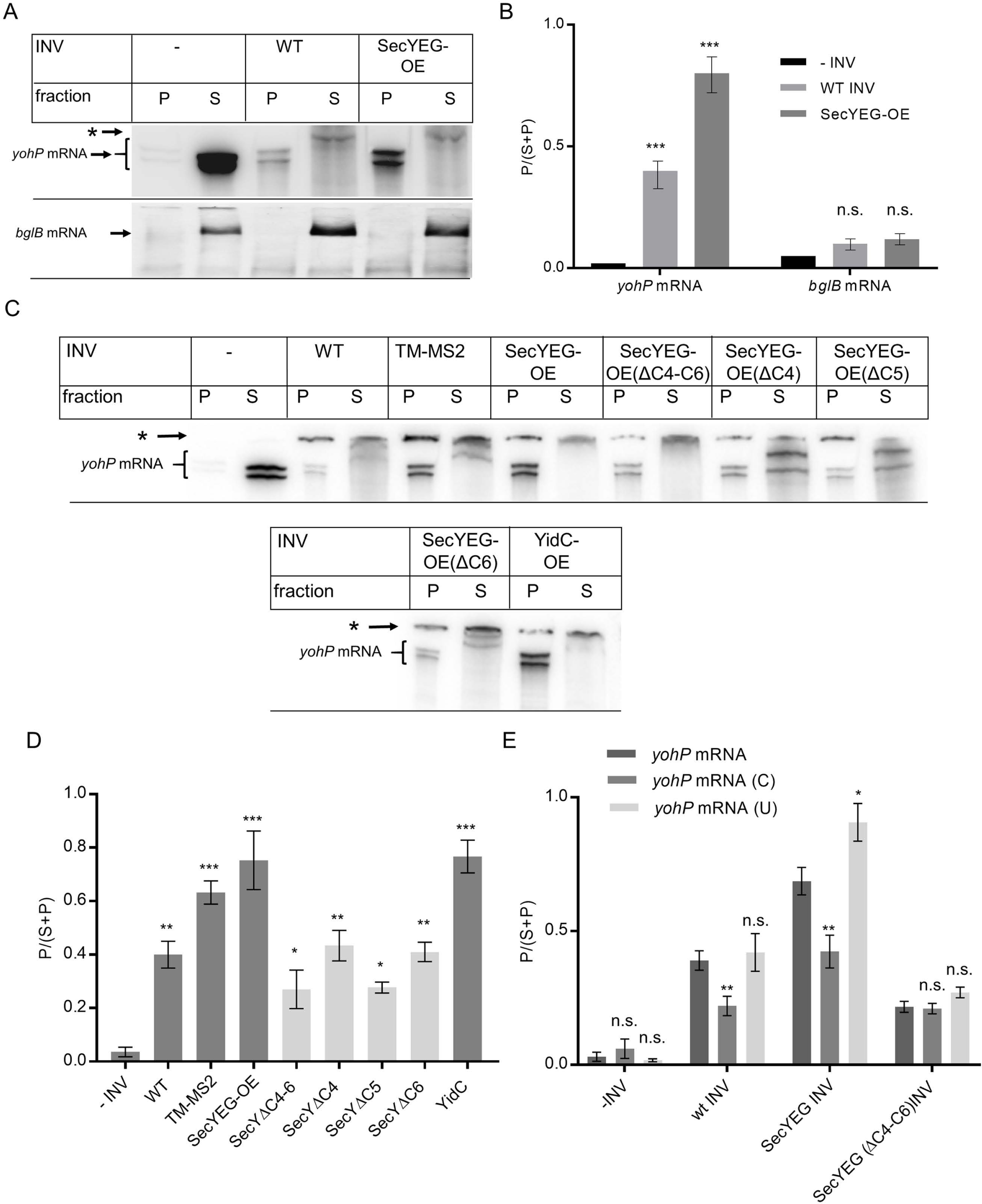
The SecYEG translocon and YidC constitute putative mRNA receptors at the E. coli membrane. **(A)** The *yohP* and *bglB* mRNAs were *in vitro* transcribed and ^32^P labeled. After purification, the mRNA was incubated with either buffer or inner membrane vesicles (INVs) isolated from either wild type *E. coli* cells (WT), or a SecYEG-overproducing strain (SecYEG-OE). After incubation, INVs and the bound mRNA were pelleted by centrifugation and the membrane fraction (P) and the soluble fraction (S) were separated on a urea-gel and analyzed by autoradiography. (See also Figure S4). **(B)** Quantification of at least three independent experiments as shown in (A) via *Imagequant*. For quantification, the radioactive signal in the pellet fraction was divided by the sum of the radioactive signals in the pellet and soluble fractions. Shown are the mean values and the standard errors of the mean (SEM), which were calculated via *GraphpadPrism.* **(C)** The mRNA binding assay was performed as in (A) using INVs from different *E. coli* strains. TM-MS2 refers to INVs from a strain over-producing a membrane-tethered MS2 protein. INVs from *E. coli* strains over-producing SecYEG variants that lacked the three cytosolic loops C4-C6 of SecY, or the individual loops were also analyzed. In addition, INVs from cells over-producing YidC were tested. (See also Figure S5). (*) corresponds to radioactive material that did not enter the gel. **(D)** Quantification of the data shown in (C) was performed as in (B). **(E)** Binding of wild type, uracil-rich (U) and cytosine-rich (C) mRNAs to the indicated INVs. Quantification of at least three independent experiments was performed as in (B). Statistical analyses were performed with the Satterthwaite corrected unpaired two-sided Student t-test, using the amount of mRNA in the pellet after incubation without INVs as reference. (*) refers to *p*-values ≤0.05; (**) to *p*-values ≤0.01, and (***) to *p*-values ≤0.001. *n.s.* denotes non-significant changes.

The eukaryotic Sec61 complex was shown to bind mRNA ^64^ and it was therefore tested whether the homologous SecYEG complex ^6^ is also involved in mRNA binding. INVs from an *E. coli* strain overproducing the SecYEG complex (SecYEG-OE INVs; Fig. S4D) showed enhanced mRNA binding (Fig. 4A, B), suggesting that the bacterial SecYEG translocon could provide a binding site for mRNA. As a control, the sedimentation assay was repeated with *bglB* mRNA, which *in vivo* is evenly distributed within the cytosol (Fig. 1A). In agreement with the *in vivo* data, the *bglB* mRNA was almost exclusively found in the supernatant after centrifugation, indicating that it does not efficiently bind to the bacterial membrane, even at higher SecYEG concentrations (Fig. 4A, B).

The increased binding of the *yohP* mRNA to SecYEG-OE INVs was further validated by a control experiment with INVs from *E. coli* cells that expressed a membrane-tethered MS2-Venus (TM-MS2) variant, which acts as a *bona fide* mRNA-binding protein. TM-MS2-Venus was constructed by fusing MS2-Venus to the first two transmembrane domains (TM) of the inner membrane protein SecY (Fig. S4E). Fluorescence microscopy confirmed the exclusive membrane localization of TM-MS2-Venus (Fig. S4E). mRNA binding to TM-MS2-Venus containing INVs was enhanced in comparison to wild type INVs and comparable to that of SecYEG-OE INVs (Fig. 4C, D), further supporting a role of the SecYEG translocon as membrane-bound mRNA receptor. SecY contains three cytosolic loops (C4-C6), which are required for binding ribosomes and targeting factors ^28, 29, 32, 67–72^. The contribution of these loops to mRNA binding was determined by constructing a SecY variant that lacked these loops (Fig. S4D). Fluorescence microscopy of a YFP-labelled SecY(ΔC4-C6)EG complex showed exclusive membrane localization *in vivo* (Fig. S5A), indicating that the truncated SecYEG complex is stably inserted into the membrane. However, the deletion of the cytosolic loops in SecY reduced membrane binding of the *yohP* mRNA (Figs. 4C & D), indicating the involvement of the C4-C6 loops in mRNA binding. This was further explored by analysing SecY variants with single loop deletions (Figs. 4C & D; Fig. S5A). Of the single loop deletions, lack of the C5-loop had the strongest effect on mRNA binding (Figs. 4C & D). The C5 loop is also the primary contact site for the ribosome ^20, 73–75^ and the SRP receptor FtsY ^29, 32^. This supports the conclusion that the SecYEG translocon can serve as mRNA receptor and that the three cytosolic loops and in particular the C5 loop of SecY likely provide mRNA binding sites. This is in line with data showing extensive contacts between the SecYEG translocon and ribosomal RNAs ^76, 77^. These experiments were performed in the presence of native SecYEG, because SecYEG is essential in *E. coli* ^6, 10^ and its conditional depletion cause dramatic secondary effects ^78–82^. The presence of endogenous SecYEG potentially explains the residual mRNA binding observed in SecY(ΔC4-C6)EG containing INVs. Still it is likely that in addition to the SecYEG translocon, other proteins can serve as membrane-bound mRNA receptors.

The YidC insertase also binds ribosomes during co-translational membrane protein insertion ^11, 83–86^. We therefore tested INVs of a YidC overproducing *E. coli* strain (Fig. S5B). These INVs also showed enhanced mRNA binding (Figs. 4C & D), which was comparable with mRNA binding to SecYEG-OE INVs. In summary, SecYEG and YidC potentially serve as mRNA receptors at the *E. coli* membrane.

The *in vivo* results show that membrane localization is determined by nucleotide composition (Fig. 2A). In support of this, the C-rich mRNA showed reduced membrane binding to wild type INVs and SecYEG-OE INVs, when compared to wild type *yohP* mRNA. In contrast, the U-rich mRNA showed enhanced binding to SecYEG-OE INVs (Fig. 4E). The cytosine and uracil content had no major influence on the residual mRNA binding to INVs from the ΔC4-C6 strain. In summary, these data demonstrate that the uracil content is an important determinant of mRNA localization both *in vivo* and *in vitro*. They furthermore suggest that uracil-rich mRNAs are preferentially bound by the SecYEG translocon.

The mRNAs tested in the *in vivo* and *in vitro* binding assays did not contain a Shine-Dalgarno sequence and the *in vitro* binding assays were performed with purified mRNAs/INVs, *i.e.* in the absence of components required for translation. This indicates that mRNA binding to the membrane is translation-independent, which was further validated *in vivo* by monitoring mRNA localization in the presence of the antibiotics kasugamycin or puromycin. However, neither antibiotic significantly influenced the localization of the *yohP*, *secY* or *bglB* mRNAs (Fig. S6), confirming their translation-independent mRNA targeting.

### Membrane-bound *yohP* mRNAs are translated and the translation product is inserted into the membrane

For analysing translation-independent mRNA targeting via the MS2 system, translation is usually blocked by antibiotics ^42, 43, 46^ (Fig. S6) or prevented by the deletion of the Shine-Dalgarno sequence (Figs 1-3). Therefore, it is unknown whether membrane-bound mRNAs are translated and whether the protein is membrane-inserted. This is particularly important because putative mRNA targeting factors, such as cold-shock proteins ^49^, can inhibit translation ^87^.

The membrane-tethered MS2 protein (TM-MS2) efficiently binds mRNA (Figs. 4 & S4) and we therefore incubated a *yohP*-MS2 mRNA variant that contained the Shine-Dalgarno sequence with TM-MS2-venus containing INVs. The INVs and the bound mRNAs were subsequently isolated by centrifugation and added to an *in vitro* translation system ^88^, containing radioactively labelled methionine and cysteine residues. This allows the detection of the *in vitro* translated YohP protein by autoradiography. Previous data had shown that membrane-inserted YohP is proteinase K (PK) protected and thus PK resistance monitors YohP membrane insertion^53^. Thus, after *in vitro* translation, one half of the sample was treated with PK to reveal membrane insertion of YohP.

In the absence of INVs, only a weak YohP band at 4.5 kDa was detectable, which likely reflects translation from *yohP* mRNAs that were pelleted even in the absence of INVs. However, YohP was largely degraded by the addition of PK (Fig. 5A). In contrast, in the presence of TM-MS2 INVs, YohP was readily detectable and almost completely protected against PK treatment (Fig. 5A). This indicates that the *yohP* mRNA binds to the TM-MS2 INVs and that the INV-bound mRNA is efficiently translated into protein, which is then inserted into the membrane. It additionally verifies that the MS2-stem loop in the *yohP* mRNA does not prevent translation and subsequent YohP insertion into the membrane (Fig. S5C).

**Figure 5:**
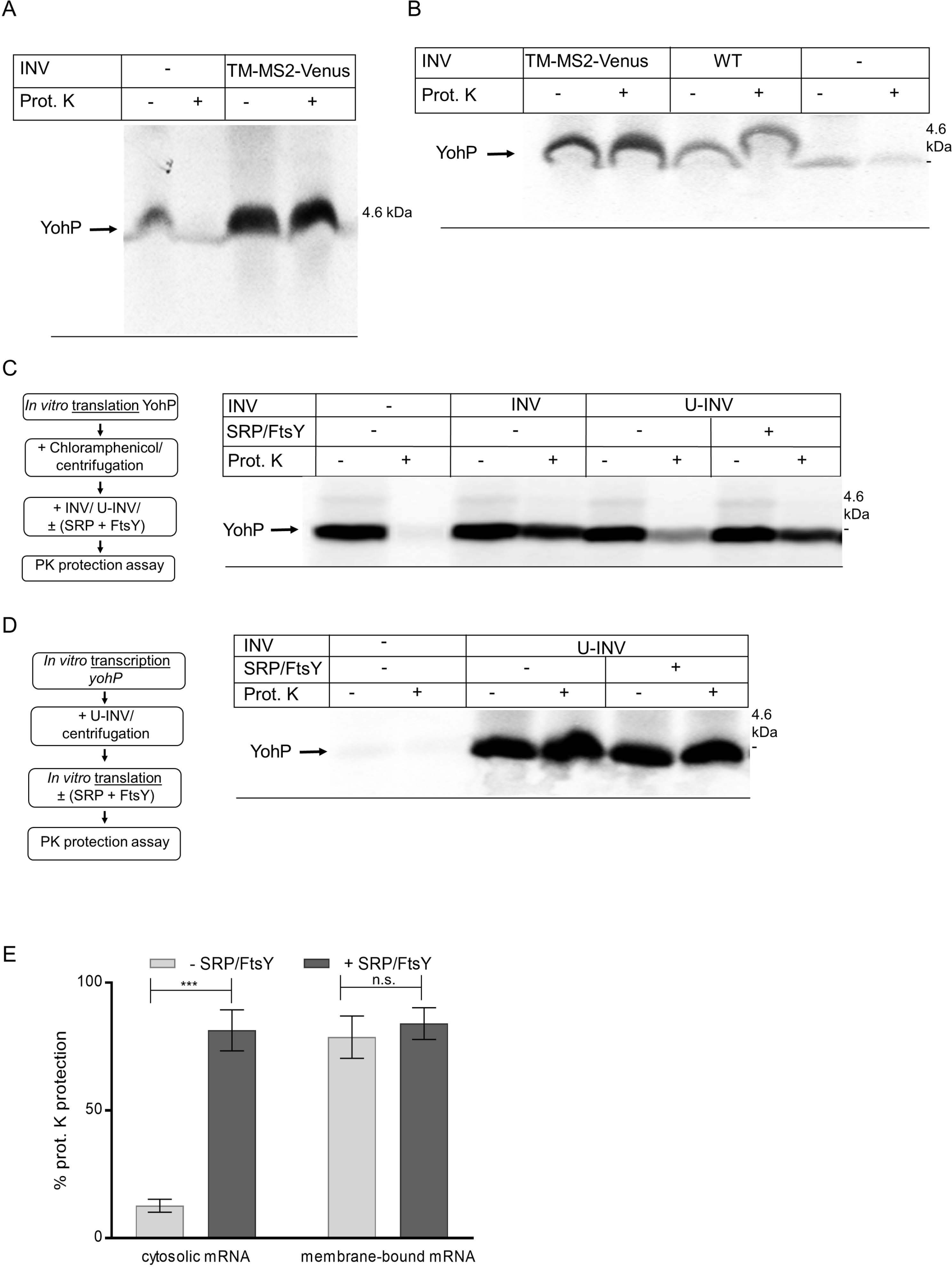
YohP translated from membrane-bound mRNAs does not require the SRP pathway for insertion. **(A)** *YohP* mRNA was incubated with TM-MS2-containing INVs or INV buffer and INV-bound mRNA was subsequently isolated by centrifugation. The pellet fraction was then incubated with an *in vitro* translation system containing ^35^S-labelled methionine and cysteine. After 30 min of incubation, proteinase K (PK) was added when indicated for monitoring YohP insertion into the membrane. **(B)** As in (A), but also with wild type INVs as further control. **(C)** YohP was synthesized from cytosolic mRNAs and post-translational membrane insertion of YohP was analyzed after *in vitro* synthesis and subsequent chloramphenicol treatment and centrifugation for stopping translation and for removing ribosomes. *In vitro* synthesized and ^35^S-labelled YohP was then incubated with INV-buffer, INVs or urea-treated INVs (U-INV). When indicated, purified SRP and FtsY (20 ng/µl) were added and membrane insertion of YohP was monitored by PK protection. (See also Figure S7). (**D**) The *yohP* mRNA was *in vitro* transcribed and purified. Purified mRNA was then added to U-INVs and U-INV-bound mRNA was re-isolated by centrifugation. The membrane-bound mRNA was translated by the *in vitro* translation system in the presence or absence of SRP/FtsY (20ng/µl, each). YohP insertion was then monitored by PK protection. (**E**) Quantification of the YohP insertion into U-INV derived from the data in (C) (YohP synthesis from cytosolic mRNAs) and (D) (YohP synthesis from membrane-bound mRNAs). Shown are the mean values of three independent experiments and the standard errors of the mean (SEM), which were calculated via *GraphpadPrism.* Statistical analyses were performed as in Fig. 4, using the YohP insertion in the absence of SRP/FtsY as reference. (***) refers to *p*-values ≤0.001 and *n.s.* denotes non-significant changes.

MS2 has a very high affinity for the MS2-stem loop ^62^ and TM-MS2 INVs bind more *yohP*-MS2 mRNA than wild type INVs (Fig. 4C). This was further verified by comparing binding of the *yohP*-MS2 mRNA to either TM-MS2-INVs or wild type INVs, followed by subsequent *in vitro* translation and PK protection. The *yohP* mRNA also binds to wild type INVs, although the amount of the translated product was lower than with TM-MS2-INVs (Fig. 5B). Thus, wild type INVs bind less mRNA than TM-MS2-INVs, but in both cases, the mRNAs were efficiently translated and the translation product was inserted into the membrane.

These data demonstrate that membrane-bound mRNAs are recognized and translated by *E. coli* ribosomes, followed by the subsequent insertion of the translation product into the membrane.

### The SRP pathway is not required for membrane insertion when YohP is translated from membrane-bound mRNAs

Targeting and insertion of bacterial membrane proteins is usually accomplished by the SRP pathway, which delivers proteins to either SecYEG or YidC for insertion ^11, 32, 89^. This raised the question of whether proteins that were translated from already membrane-bound mRNAs would still require the SRP pathway for insertion. This was tested *in vitro* by employing urea-treated INVs (U-INVs). Urea-treatment removes most SRP and the SRP receptor FtsY from INVs (Fig. S7A) and prevents efficient membrane protein insertion, unless purified SRP and FtsY are added ^88, 90^. In a first experiment, YohP was *in vitro* synthesized, translation was stopped by the addition of chloramphenicol and the *in vitro* synthesized YohP was incubated with INVs or U-INVs, supplemented with SRP/FtsY when indicated. In the absence of INVs, YohP was completely degraded, while in the presence of INVs, YohP was largely protease resistant, indicating its membrane insertion (Fig. 5C). When YohP was incubated with U-INVs, only a small portion of YohP was membrane inserted, in line with the reduced amounts of SRP/FtsY in U-INVs. However, when YohP was incubated with U-INVs in the presence of purified SRP/FtsY, YohP was efficiently inserted into the membrane. This validates that when YohP is produced from cytosolic mRNAs, it requires the SRP pathway for targeting and insertion, which supports previous studies^53, 91^.

Next, the SRP-dependency of YohP insertion was tested for YohP that was translated from already membrane-bound mRNAs. *YohP* mRNA was *in vitro* transcribed, incubated with either no INVs (buffer control) or U-INVs (Fig. 5D). The pellet after centrifugation containing the U-INV-bound mRNAs was then used for *in vitro* translation combined with a PK protection assay in the presence or absence of SRP/FtsY. Translation of the *yohP* mRNA was detected in the presence of U-INVs, but not in the buffer control. Importantly, PK protection was already observed in the absence of SRP/FtsY and not further stimulated by their presence (Fig. 5D & E). This demonstrates that membrane insertion of YohP is largely independent of the SRP pathway when YohP is translated from already membrane-bound mRNAs. Thus, bacteria can evidently bypass the SRP pathway for membrane protein insertion by translation-independent mRNA targeting.

### mRNA targeting ensures membrane protein insertion during stress conditions

The physiological advantage of employing an additional mRNA targeting strategy for YohP insertion in addition to the established signal sequence dependent targeting via the SRP pathway ^53^ is not immediately obvious. mRNA targeting could serve as a back-up strategy when the low-abundant SRP pathway is saturated ^92^ or inhibited by the stress-induced alarmones (p)ppGpp ^5, 91^. To test this latter case, we analysed whether mRNAs that show impaired membrane targeting also show reduced YohP insertion into the membrane when exposed to stress conditions. We designed an *in vivo* pulse-chase experiments to monitor membrane insertion of YohP that was translated from either wild type or the U- and C-rich mRNAs. The mRNAs were transcribed from a *T7*-dependent promoter in wild type cells supplemented with radioactively labelled methionine/cysteine. Membrane insertion of ^35^S-labelled YohP was then determined by cell fractionation after 5 min labelling (pulse) and up to ten minutes growth in the absence of radioactive amino acids (chase). Wild type YohP was detectable in the membrane fraction (P) after a 1 min chase, and then decreased gradually, presumably due to proteolysis (Fig. 6A). A similar pattern was observed for the U-rich and C-rich variants, respectively (Fig. 6A), although the C-rich *yohP* mRNA showed impaired membrane binding *in vivo* and *in vitro* (Figs. 2 & 4). Apparently, YohP is inserted into the membrane even when membrane targeting of its mRNA is compromised, likely because signal-sequence dependent targeting via the SRP pathway is still sufficient for YohP insertion. The cytosolic protein YchF served as a control for the fractionation procedure and was almost exclusively detected in the soluble fraction (Fig. S7B)

**Figure 6:**
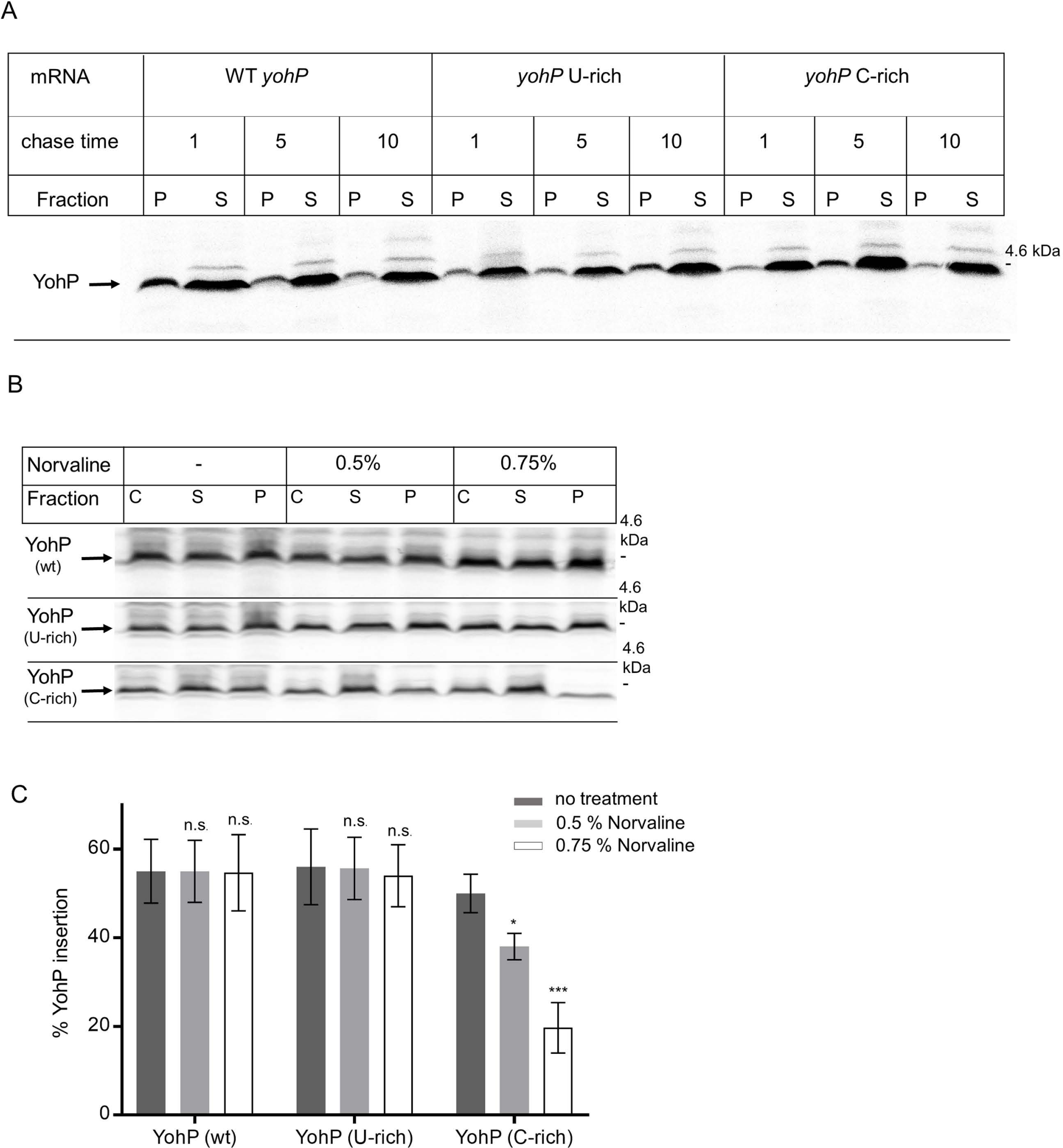
mRNA targeting and signal sequence dependent targeting via the SRP pathway act in parallel. **(A)** *In vivo* pulse chase experiments for monitoring membrane insertion of YohP. YohP variants with identical amino acid sequence were produced from three different *T7*-dependent expression plasmids, containing either wild type *yohP*, the uracil-rich or the cytosine-rich sequences (see also Figure 2 & Figure S2). After induction of *T7*-dependent mRNA production, endogenous *E. coli* RNA polymerase was blocked by rifampicin and cells were labeled for 5 min with ^35^S-labelled methionine/cysteine and then chased with non-radioactive methionine/cysteine for 1, 5 or 10 min. Cells were subsequently lysed by ultrasonic treatment and separated into the membrane fraction (P) and the cytosolic fraction (S) by ultracentrifugation. Samples were then separated by SDS-PAGE and visualized by autoradiography. **(B)** As in (A), but cells were treated when indicated with norvaline for 1.5 h before mRNA production and rifampicin treatment. Cells were 5 min pulsed and 5 min chased before cell lysis, centrifugation and further processing as in (A). (see also Figure S7). (**C**) Quantification of three independent experiments via *Imagequant*. For quantification, the radioactive signal in the pellet fraction was divided by the sum of the radioactive signals in the pellet and soluble fraction and is displayed as % YohP insertion. Shown are the mean values and the standard errors of the mean (SEM), which were calculated via *GraphpadPrism.* Statistical analyses were performed as in Fig. 4, using YohP insertion in the absence of norvaline as reference. (*) refers to *p*-values ≤0.05; (***) refers to *p*-values ≤0.001 and *n.s.* denotes non-significant changes.

The experiment was then repeated in the presence of norvaline, an amino acid derivative that induces leucine starvation and activates (p)ppGpp formation ^93–95^. (p)ppGpp accumulation inhibits SRP-dependent YohP insertion by preventing SRP-FtsY complex formation ^91^. In addition, (p)ppGpp accumulation allosterically inhibits RNA polymerase ^96^. This is visible by a significantly reduced global protein synthesis in cells treated with 0.5% norvaline (Fig. S7C). On the other hand, YohP synthesis was not drastically impaired, because *T7* RNA polymerase is insensitive to (p)ppGpp accumulation ^97^. This allowed us to determine YohP insertion in the presence of norvaline. The addition of 0.5% or 0.75% norvaline did not significantly influence the insertion of YohP that was translated from wild type *yohP* mRNA, because mRNA targeting can apparently compensate for impaired SRP-dependent targeting (Fig. 6B & C). Norvaline did also not significantly reduce the insertion of YohP that was translated from U-rich mRNAs. In contrast, the insertion of YohP that was translated from C-rich mRNAs, showed a dosage-dependent reduction in membrane insertion (Fig. 6B & C). This demonstrates that the simultaneous inhibition of SRP-dependent targeting and mRNA targeting reduces YohP membrane insertion. A complete inhibition of YohP insertion could not be accomplished, because higher norvaline concentrations prevented protein synthesis completely, likely via the competitive inhibition of GTP-dependent translation factors by (p)ppGpp ^96^.

In summary, our *in vitro* and *in vivo* data demonstrate that mRNA targeting serves as an alternative strategy for membrane protein insertion. This is likely particularly important for membrane proteins such as YohP and other small membrane proteins, which are upregulated during stationary phase or when cells encounter stress, conditions which are associated with (p)ppGpp accumulation.

## Discussion

Non-random localization of mRNAs has been demonstrated in bacteria, but the mechanisms that trigger mRNA localization are largely unknown. Equally unknown is how mRNA targeting contributes to protein transport into and across the bacterial membrane. By combining *in vivo* imaging with *in vitro* biochemical assays, our study reveals several important aspects of mRNA targeting in bacteria, using the *yohP* mRNA as model: (1) The coding sequence of the mRNA contains sufficient information for membrane targeting and does not depend on information within the 5′- or 3′UTRs or the translation product. (2) The uracil content of mRNAs is an important determinant for membrane binding of mRNAs *in vivo* and *in vitro*. (3) Increasing the cytosine or adenine content reduces membrane binding of mRNA, while increasing the guanine content does not affect the mRNA-membrane interaction. However, the influence of the nucleotide composition on membrane targeting is largely dependent on whether secondary structures within the mRNA are maintained. (4) mRNA binding to isolated bacterial membranes occurs independently of soluble targeting factors. (5) The SecYEG translocon and the YidC insertase constitute potential mRNA receptors. (6) Membrane-bound mRNAs are efficiently translated and the translation product is inserted into the membrane independently of the SRP pathway. (7) Impairment of mRNA targeting does not significantly reduce protein insertion *in vivo*, unless the SRP pathway is inhibited in parallel. The overall data suggest that mRNA targeting acts in parallel to the canonical signal sequence based targeting and ensures membrane protein insertion when the SRP pathway is saturated or inhibited.

SRP-dependent protein targeting is generally considered to be essential ^13^, but eukaryotic and bacterial cells have developed strategies to cope with impaired SRP-dependent targeting. Down-regulation of protein synthesis and up-regulation of chaperones and proteases enables yeast cells to grow when SRP is inactivated ^98^. A similar response is observed in *E. coli* ^99^, which is further amended by reducing translational speed and fidelity ^78, 100^ and by increased translation of leader-less mRNAs encoding for stress-responsive proteins ^101, 102^. In *Streptococcus mutans*, the SRP pathway can be completely inactivated due to the presence of a YidC variant with strong affinity for ribosomes and RNCs ^103, 104^. Some bacteria, such as *Leptospira sp,* lack genes for the SRP components completely ^105^, indicating that alternative pathways for membrane protein insertion exist.

In *E. coli*, there is so far no indication that SRP- and FtsY-levels are transcriptionally regulated ^5^, although SRP is post-translationally regulated via proteolysis ^26, 106^. However, a recent study has demonstrated that under stress conditions, the hyper-phosphorylated guanine nucleotides ppGpp and pppGpp bind to the GTPase domains of both SRP and FtsY and prevent the formation of a functional targeting complex ^91^. This is intriguing, because many small membrane proteins, such as YohP, are upregulated when cells encounter stress conditions ^107–110^, but the simultaneous (p)ppGpp accumulation would prevent their membrane insertion by the SRP-pathway ^96^. mRNA targeting and SRP-independent protein insertion, as shown here for YohP, could provide a solution to this conundrum and allow for the insertion of stress-response proteins even when the SRP pathway is impaired (Fig. 7). The observation that SecYEG and YidC provide binding sites for mRNA supports this hypothesis, because the primary function of the SRP pathway is to direct RNCs to either SecYEG or YidC. mRNAs that are already bound to either SecY or YidC on the other hand can engage ribosomes without prior need for a ribosome-targeting step. Binding of mRNAs to the homologous Sec61α has been shown previously ^64^ and is in line with the ability of both SecY and YidC to interact with ribosomal RNAs. Cryo-EM studies have shown that the cytoplasmic loops C4 and C5 of SecY are in contact with the 23S rRNA helices H50-H53-H59 and H6-H24-H50, respectively ^76^, which are both essential for SecY binding ^111^. The 23S rRNA consists of 2905 nucleotides with a uracil content of just 20%. However, with the exception of H24, the uracil content in the helices interacting with SecY is higher and reaches 37.5% in H53. Thus, the observed preference of SecY for uracil-rich sequences is supported by the available ribosome-SecY structures. The 23S rRNA helix 59 also contacts the cytosolic loops of YidC ^85, 112, 113^, further supporting a role of YidC as RNA receptor.

**Figure 7:**
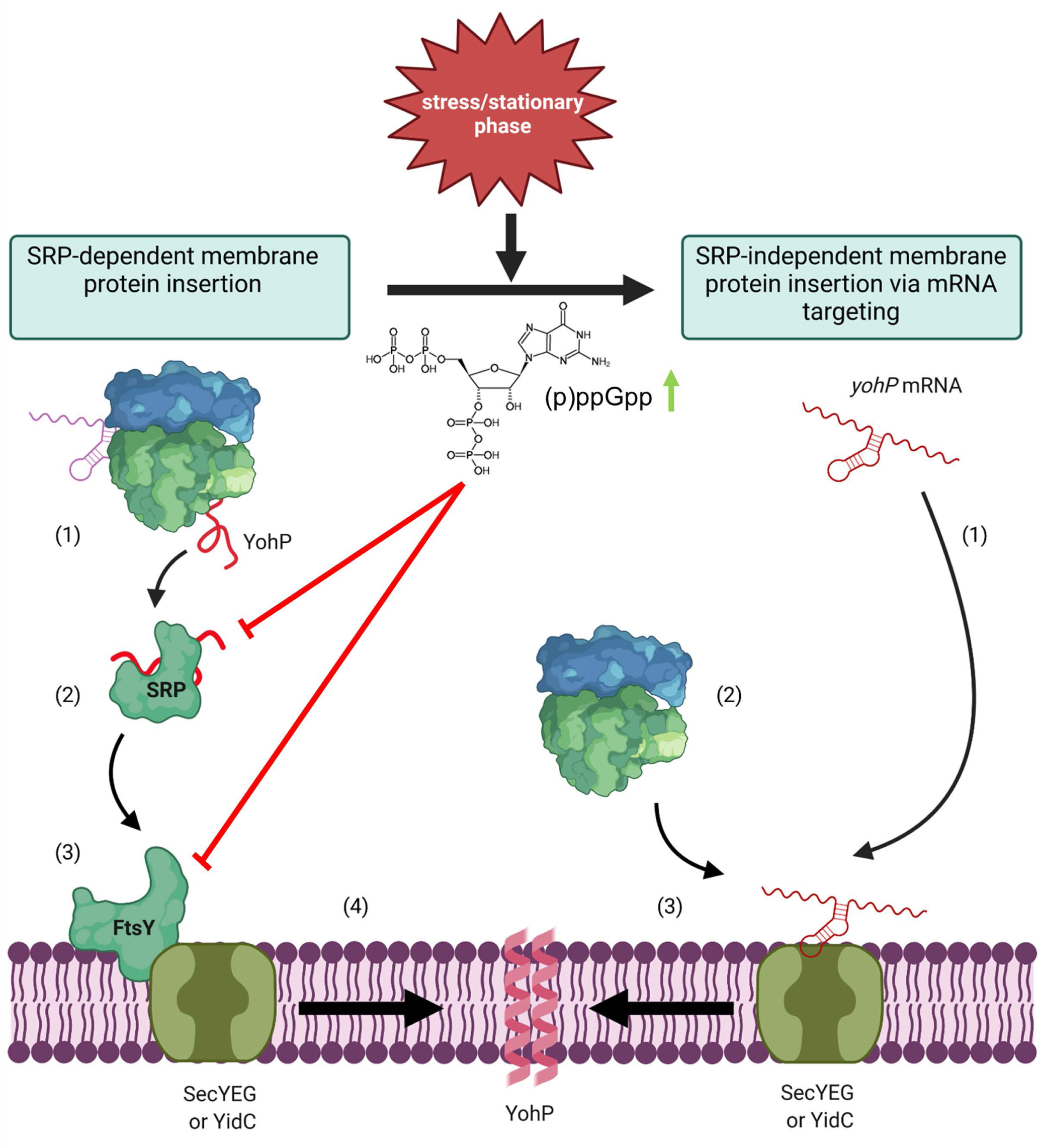
mRNA-targeting as SRP-independent back-up strategy for membrane protein insertion in *E. coli.* **Left panel:** Membrane protein insertion in *E. coli* depends on SRP and its receptor FtsY, which target client proteins to either the SecYEG translocon or the YidC insertase for insertion. **(1)** In most cases, SRP targets its substrates co-translationally, but can also act post-translationally for small membrane proteins, such as YohP ^53, 107^. **(2)** SRP targets proteins to the SRP receptor FtsY, which is associated with either the SecYEG translocon or the YidC insertase. **(3)** Subsequently, YohP is inserted into the membrane and dimerizes **(4)**. **Right panel:** During stress conditions or when cells enter stationary phase, the concentration of the alarmones (p)ppGpp increase, resulting in a cellular re-programming, which includes the inhibition of the SRP pathway by preventing SRP-FtsY complex formation. Under those conditions, mRNAs can bind directly to the SecYEG translocon or YidC **(1)** and ribosomes translate these membrane-bound mRNAs **(2)**, allowing for SRP-independent membrane insertion **(3)**. This SRP-independent pathway is likely particularly important for membrane proteins, which are up-regulated during stress conditions or when cells enter stationary phase, such as YohP. The cartoon was generated using Biorender (https://biorender.com/).

It is generally assumed that transcription and translation are coupled in bacteria ^114–116^, but the small size of the *yohP* mRNA potentially prevents such coupling. Still, translation-independent mRNA targeting has also been shown for larger transcripts ^42, 43, 117^, demonstrating that uncoupled transcription-translation is not a particular feature of small transcripts. Nevertheless, we can currently not exclude that the insertion of proteins translated from larger membrane-bound mRNAs or at least some of them would still depend on SRP. This would explain why bacterial cells survive SRP depletion for some time, but ultimately loose viability. Still, our data indicate that membrane targeting of mRNAs provides a back-up strategy and complements the canonical signal-sequence dependent targeting, in particular when SRP-dependent targeting is compromised.

## Acknowledgments

The authors thank Veronika Erichsen and Rosmery Rupp-Falconi for assistance and Werner Bigott for help with Northern blot experiments. HGK acknowledges support from the Deutsche Forschungsgemeinschaft (DFG) (grants KO2184/8, KO2184/9 (SPP2002); SFB1381, Project-ID 403222702, and RTG 2202, Project-ID 278002225). AW is supported by the DFG with grants SFB1381, Project-ID 403222702 and GRK2344/322977937. The Kraft laboratory has received funding from the DFG, Project ID 409673687, SFB 1381 (Project ID 403222702), SFB 1177 (Project ID 259130777), from the European Research Council (ERC) under the European Union’s Horizon 2020 research and innovation program under grant agreement No 769065, and from the European Union’s Horizon 2020 research and innovation program under grant agreement No 765912. This work reflects only the authors’ view and the European Union’s Horizon 2020 research and innovation program is not responsible for any use that may be made of the information it contains. The authors declare no competing financial interests. WS acknowledges support from the Traditional Chinese Hospital of Tongshan Xuzhou, China and the Chinese Scholarship Council. Work included in this study has been performed in partial fulfillment of the requirements for the doctoral theses of PS, WS, EL, ML at the University of Freiburg.

## Author contributions

Conceptualization: HGK, MHU, AW. Investigation: PS, AO, WS, MHU, EL, ML. Visualization: PS, WS, MHU, AW, HGK; Funding acquisition: HGK, AW, CK; Supervision: HGK, MHU, AW; CK Writing: PS, WS, MHU, AW, CK, HGK. All authors have read and commented on the manuscript.

## Data availability

All data are present in the current manuscript.

## Resource availability

### Lead contact

Further information and requests for resources and reagents should be directed to the lead contact, Hans-Georg Koch

### Materials availability

All plasmids are available upon request, subject to a material transfer agreement (MTA), from Hans-Georg Koch

### Data availability

All data are present in the current manuscript.

### Experimental model and subject details

All bacterial strains used in this study are derived from wild type *E. coli*K-12 strain and are listed in supplementary table S1. Plasmids and oligonucleotides used in this study are listed in the supplementary tables S2 and S3, respectively. All plasmids were constructed by using the Gibson assembly protocol using 100 ng vector and a 1:3 vector /insert ratio and the NEB Site directed mutagenesis kit using the manufacture’s protocol. For generating the A-rich YohP sequence, oligonucleotides containing the entire sequence were used and the construct was assembled by using the HiFi DNA assembly Master Mix (NEB). The G-rich YohP sequence was first cloned into plasmid pUC57-BsaI-Free by BioCat GmbH (Heidelberg, Germany) and then re-cloned into the pSC_MS2.6x plasmid via Gibson assembly.

## Method details

### Delta Vision^TM^ Ultra microscopy

mRNA localization was monitored in *E. coli* BL21 cells carrying the plasmid pBAD24-MS2-Venus and the vector pSC.YohP-MS2.6x. The latter plasmid contained the *yohP* sequence fused to the MS2-recognition sequence, but lacked a Shine-Dalgarno sequence. Cells were inoculated overnight with 50 µg/mL ampicillin and 35 µg/mL chloramphenicol. The following day a pre-culture was inoculated 1:100 and grown until an OD_600_ = 0.5, cells were then induced with 1 mM IPTG for 1h to induce mRNA expression, and with 0.2 mM arabinose to induce the MS2-GFP protein for 1.5h. 1 mL cell culture (corresponding to 2 x 10^8^ cells) was centrifuged for 10 min at 2,300 x g at room temperature, and the pellet was after washing re-suspended in 800 µL PBS followed by Nile Red staining (0.0035 g Nile Red was dissolved in 1 ml acetone and diluted 1:20 in the bacterial cell suspension). The cell suspension was then incubated at room temperature for 20 min. After staining, 10 µl cell culture was transferred to a clean Glass Bottom Dish (35 mm Dish with 20 mm Bottom Well # 1.5 Glass (0.16-0.19 mm)) (Cellvis, Mountain View, USA) and then covered with 800 µL of PBS/1 % agarose and a cover slip. During imaging, the background fluorescence from *E. coli* cells was also monitored by growing cells under the same conditions but without any inducer and by analyzing plasmid-free *E. coli* strains.

The same imaging conditions were used for monitoring protein and mRNA localization. Imaging was performed with a Delta Vision^TM^ Ultra High Resolution Widefield Microscope using a triple beamspliter emission filter set (395/495/610) (GE Healthcare, Munich, Germany) at 100x magnification and a numerical aperture of 1.35 with an Olympus UPlanSapo objective (Olympus, Hamburg Germany) and immersion oil with a refractive index of 1.518. Images were taken at room temperature. Exposure time was 0.4 s for Venus and at 32 % laser power. Recording, using camera sCMOS pro edge (PCO, Kelheim, Germany), was performed using a 3 µm Z-scan with optical section spacing of 0.1 µm. Acquired images were deconvolved using the *Acquire ultra*-software (softWoRx, GE Healthcare, Munich, Germany) and further analysed with *ImageJ/ Fiji*.

### Fluorescence in situ hybridization

For visualization of *yohP* RNA, a set of 19 DNA oligonucleotide probes (supplementary table S4) was designed using the Stellaris RNA-FISH Probe Designer program (https://www.biosearchtech.com/stellaris-designer). Each probe against the *yohP* RNA (excluding the MS2 tag) was created with a GC content of approximately 50%, (18 nt each, masking parameter: 0) and 3’ labeled with TAMRA fluorophore (5-carboxytetramethylrhodamine). Plasmids pSc containing *yohP* with or without MS2-*6xbs* were grown until OD_600_ 0.5 and mRNA expression was induced for 1 h with IPTG. Wild type *E.coli* BL21 cells for controls (- probe; *psbA* probe) were grown under identical conditions, but without IPTG addition. Cell fixation and *in situ* hybridization was performed as described by Skinner *et al.* ^118^. 5 ul of the cell suspension was transferred to a Glass Bottom Dish with the same specifications as described for mRNA localization. Imaging was performed using the same conditions that were used for mRNA localization except that the exposure time was set to 0.1 s with 100% laser intensity.

### In vitro transcription

For *in vitro* transcription of *yohP,* pRS1 containing the YohP-MS2-6xbs coding sequence was first linearized by Eco*RI* restriction digest (NEB, Frankfurt, Germany). *In vitro* transcription was performed using AmpliScribe T7-Flash Transcription kit (Lucigen Corp., Middleton, USA), in the presence of ^32^P-labeled nucleotides. The RNA were purified by using the *RNA Purification Kit* (Qiagen, Hilden, Germany) and stored at –80 °C. For *in vitro* transcription of the 4.5S RNA, the plasmid pT7/3a coding for 4.5S RNA was linearized using Bam*HI* (NEB) and *in vitro* transcription was performed as above, but without radioactively labelled nucleotides.

### Total RNA isolation and Northern blot hybridization

Total RNA was isolated from *E. coli* cells via the PGTX protocol ^119^ and RNA quality was controlled by agarose gel electrophoresis. For Northern blot analyses, 2 µg of total RNA was separated on a 1.3% agarose gel and blotted onto a nylon membrane by capillary blotting in 10-fold SSC buffer (1.5 M NaCl, 150 mM trisodium citrate, pH 7.0). After blotting the RNA was cross-linked to the nylon membrane for 12 s at 100 µJ/cm^2^. As a probe for MS2-stem-loop containing RNAs, the oligonucleotide 5′-CTGCAGACATGGGTGATCCTCATGT-3‵was labelled with ^32^P (30 µCi γ-^32^P-ATP) using T4-polynucleotide kinase and added to the hybridization buffer (120 mM Na_3_-PO_4_, 250 mM NaCl, 7% SDS, pH 7.2). Hybridization was performed overnight at 45 °C and the membrane was washed sequentially by pre-heated buffer III (2 x SSC, 1% SDS) and pre-heated buffer IV (1 x SSC, 0.5 % SDS). The radioactive signal on the membrane was then detected by phosphoimaging (Typhoon FLA9500 imaging system, GE Healthcare, USA).

### *In vitro* protein synthesis, proteinase K protection assays and protein purification

For protein transport assays, proteins were synthesized *in vitro* using a purified transcription/translation system composed of cytosolic translation factors (CTF) and high salt washed ribosomes ^88^. The ^35^S-Methionine labeling mix was obtained from Hartmann Analytic (Braunschweig, Germany). Inverted inner membrane vesicles (INVs) of *E. coli* cells were prepared by growing *E. coli*cells to approx. OD_600_= 1.2 on LB medium. Cells were harvested and resuspended in INV buffer (50 mM triethanolamine acetate, pH 7.5, 200 mM sucrose, 1 mM DTT). Next, the samples were lysed in the presence of protease inhibitors by French pressing (*Thermo scientific*, Langenselbold, Germany) at 800 psi and the cell debris was removed by centrifugation at 30,000g for 30 minutes in a SS34 rotor. The supernatant (S30) was further centrifuged at 150,000g for 2 hours in a TLA 50.2 rotor and the pellet containing the crude bacterial membranes was dissolved in INV buffer, loaded onto a step-wise sucrose gradient (0.77 M, 1.44 M and 2.02 M sucrose in INV buffer) and the inner membrane fraction (inverted inner membrane vesicles, INVs) and the outer membrane fraction were separated as described ^120^. The INV fraction was diluted four-fold in INV buffer and centrifuged at 150,000g for 2 hours in a TLA 50.2 rotor. The pellet was then resuspended in INV buffer. Urea-treated vesicles (U-INV) were generated by incubating INVs for 1 h with 6 M urea in INV buffer on ice. Subsequently, U-INVs were diluted four-fold with INV buffer and centrifuged in a TLA55 rotor for 2 h at 55,000 rpm through a 750 mM sucrose cushion in INV buffer. The pellet was resuspended in INV buffer and centrifuged again as above. After the second centrifugation, the pellet was resuspended in INV buffer and stored at -80 °C.

Membrane insertion of YohP was determined by proteinase K protection assays. After *in vitro* synthesis of YohP, samples were incubated for 10 min at 37 °C with 35 µg/ml chloramphenicol for inhibiting translation and then centrifuged for 30 min at 55,000 rpm in a Beckmann TLA55 rotor. The supernatant containing *in vitro* synthesized YohP was then incubated with INV or U-INV (11 mg protein/mL) for 10 min at 37 °C, in the presence or absence of SRP and FtsY (20 ng/µL each). One half of the reaction was directly precipitated with 10% trichloroacetic acid (TCA), while the other half was first treated with 0.5 mg/mL proteinase K for 20 min at 25 °C and only then TCA precipitated. Proteinase K was inactivated in 10% TCA by incubation for 10 min at 56 °C. Next the samples were denatured at 56 °C for 10 minutes in 35 μL of TCA loading dye (prepared by mixing one part of Solution III (1M dithiothreitol) with 4 parts of Solution II (6.33% SDS (w/v), 0.083M Tris-Base, 30% glycerol and 0.053% Bromophenol blue) and 5 parts of Solution I (0.2M Tris, 0.02M EDTA pH 8)) and separated on SDS Page before phosphorimaging.

For *in vitro* translation of membrane-bound mRNAs, *in vitro* transcribed and purified mRNAs (2 µg/µL) were incubated with INVs or U-INV (11 mg/mL) for 5 min at 4 °C. Samples were then centrifuged for 40 min at 150.000g in a Beckmann TLA100.3 rotor. The pellet was resuspended in INV buffer and added to the *in vitro* translation system. When indicated, purified SRP and FtsY were present during this incubation (20 ng/µL each). Subsequently, membrane insertion of YohP was analyzed by proteinase K protection assays, as described above.

Protein purification followed previously described protocols for Ffh ^121^ and FtsY ^23^. Ffh was concentrated on a 10 kDa centrifugal filter (Amicon Ultra, Witten, Germany) and re-buffered in HT buffer + 50% Glycerol (50 mM HEPES, pH 7.6, 100 mM KOAc, pH 7.5, 10 mM Mg(OAc)2, 1mM DTT) using a PD-10 column (GE Healthcare, Munich Germany). The protein was stored at -20 °C. SRP was reconstituted by incubating 0.1 mg mL^-1^ *in vitro* transcribed 4.5*S* RNA with 1.5 µM Ffh for 15 min at 25 °C in HT buffer. FtsY was re-buffered in HT buffer using a PD-10 Column (GE Healthcare, Munich, Germany) and stored at -80 °C.

### Separation of small membrane proteins by SDS-PAGE and western blotting

For separation of small membrane proteins, a modified Tris-Tricine-SDS-PAGE system was used ^122^. Gels were casted and gel electrophoresis was performed in a vertical dual gel system (Peqlab, Erlangen, Germany) with constant cooling to 3 °C. Small membrane proteins were separated by 16.5% Tricine-SDS gels with an acrylamide/ bis-acrylamide 48:1.5 or 37.5:1 ratio for *in vitro* expressed proteins. Gel electrophoresis was performed overnight at 4°C and 25-27 mA. If gel drying was required for subsequent autoradiography, gels were fixed for 30-60 minutes in 35 % ethanol and 15 % acetic acid and washed three times with distilled water for 15 minutes each.

### Immune detection and antibodies

For immune detection of small proteins after SDS-PAGE, samples were electroblotted onto PSQ 0.2 µm membranes (GE Healthcare, Munich, Germany) with 2 mA/cm^2^ in a semi-dry system (transfer buffer 48 mM Tris, 39 mM Glycine, 20 % methanol (v/v)). Larger proteins were electroblotted onto Nitrocellulose 0.45 µm membranes (GE Healthcare) with 750 mA for two hours in a tank buffer system (transfer-buffer: 50 mM Tris, 384 mM Glycine, 20 % Ethanol (v/v), 0.02 % SDS (w/v)). Membranes were blocked with 5 % milk powder in T-TBS buffer for at least 1 h. Polyclonal antibodies against the complete and SDS-denatured proteins Ffh, FtsY and YidC were raised in rabbits ^88^. Antibodies against the SecY peptide MAKQPGLDFQSAKGGLGELKRRC were raised in rabbits by GenScript Biotech (Leiden, Netherlands) and have been validated before ^30, 121, 123^. Monoclonal peroxidase-conjugated antibodies against the His6-tag (HisProbe™-HRP Conjugate) were purchased from Thermo Scientific (Langenselbold, Germany) and from Roche (Grenzach-Whylen, Germany). Peroxidase-coupled goat anti-rabbit antibodies (Caltag Laboratories, Burlingham, CA, USA) were used as secondary antibodies with ECL (GE Healthcare, Munich, Germany).

### Preparation of liposomes from synthetic phospholipids

Synthetic Lipids (PE (1,2-Dioleyl-sn-Glycero-3-Phosphoethanolamine)), PG (1-Palmitoyl-2-Oleoyl-sn-Glycero-3-(Phospho-rac-1-Glycerol)), CL (1,1’2,2’-Tetraoleoyl-Cardiolipin) PC (1,2-Dioleoyl-sn-Glycero-3-Phosphocholine) were purchased from Avanti Polar Lipids (Alabaster, AL, USA). Lipids were withdrawn from their flasks under an argon atmosphere using a gas-tight Hamilton syringe. A dry lipid film was generated by evaporating the sample in a speed-vac for 1h at 25°C. Hydration of lipids was accomplished by adding INV-buffer to obtain a final concentration of 2.5 mg/mL. Disruption of LMV (large multilamellar vesicles) occurred in a bath sonicator for 10 min, followed by repeated freezing and thawing cycles.

### Membrane binding assay for mRNA

For membrane binding assays, all buffers were prepared with DEPC-treated water for reducing RNase activity. INV (40 µg) or liposomes (2.5 µg) were incubated with 2 uL of the RNase inhibitor RNasin for 10 min on ice, before 0.5 μL of ^32^P-labelled mRNA was added. The reaction was adjusted to 20 μL with INV buffer and further incubated at 4 °C for 5 min. Samples were centrifuged at 150.000g in a Beckmann TLA100.3 rotor for 1 h at 4 ℃. After centrifugation, the supernatant and the pellet fractions were withdrawn and 20 µL RNA loading dye was added (95 % formamide, 0.025 % SDS, 0.1 % bromophenol blue, 0.5 mM EDTA). Samples were denatured at 90 °C for 5 min and separated on a urea-gel (24 g urea, 12.5 mL 40 % acrylamide-bis acrylamide (29:1), 166 μL 30 % APS, 20 μL TEMED, 5 mL 10 x TBE, adjusted to a final volume of 50 ml with water). The gel was pre-run at 260V for 1h and then the wells were rinsed with running buffer (1 x TBE). Samples were separated at 260V until the marker dye front reached the end of the gel. After fixing the gel (1x TBE plus 10 % methanol and ethanol) for 10min, the gel was dried and analyzed by phosphorimaging.

### *In vivo* pulse labeling followed by cell fractionation

10 mL LB medium were inoculated with BL21 cells carrying pRS1-YohP (WT) or its U-rich and C-rich variants. 100 μg/μL of ampicillin were used for all the constructs and cells were grown overnight at 37°C. 200 µL of these cultures were then harvested, washed twice with M63 medium (20 g/L glycerol, 13.6 g/L KH_2_PO_4_, 2 g/L [NH_4_]_2_SO_4_, 0.5 mg/L FeSO_4_ [pH 7.0] adjusted with KOH, 0.5 mg/mL thiamine, and 0.1 mM of 18 amino acids mix [all amino acids with exception of cysteine and methionine]) and resuspended in 20 mL of M63 medium, supplemented with 100 μg/μL ampicillin. The cultures were grown at 37 °C until O.D 0.5-0.6, and protein production was induced with 0.5 mM IPTG followed by incubation for at least 1 h. When indicated, norvaline was added together with IPTG to a final concentration of 0.5 or 0.75%. After 1 - 1.5h of growth, 50 μg/mL of rifampicin were added to block the endogenous RNA polymerase and samples were further incubated for 30 min. Subsequently, 15 × 10^8^ cells were collected and transferred to 2 mL fresh M63 media. Samples were pulsed for 5 minutes at 37 °C by adding 2 mL of ^35^S-L-methionine and ^35^S-L-cysteine labeling mix (11 mCi/mL, Hartmann Analytics; Braunschweig, Germany). Samples were chased by adding non-radioactive methionine and cysteine (1 µg each) to each samples. 250 μL of the samples were collected from each tube after 1 min, 5 min, 10 min chase. Samples were harvested, resuspended in 1 ml lysis buffer (50 mM Tris/HCl, pH 7.5; 50 mM NaCl, 1 mM EDTA, 1 mM DTT) and lysed by ultrasonic treatment (five 15 second pulses on ice) in the presence of PMSF (0.1 mM). Unbroken cells were pelleted by a 15 min centrifugation in an Eppendorf FA-45-30-11 rotor at 30,000g and 4 °C. The supernatant was further centrifuged in a Beckmann TLA 100.3 rotor at 120,000g for 1 hour at 4 °C to pellet the bacterial membranes. The pellet was directly denatured with 30 μL protein loading dye at 45 °C for 20 minutes. The supernatant was first precipitated with trichloracetic acid (5 % final concentration), and then denatured. All samples were separated by 16 % Tricine-SDS-PAGE and analyzed by phosphorimaging.

### Quantification and statistical analyses

The brightness and contrast of the images were adjusted with *Fiji/ImageJ* and further processed by a multi-profile measurement plug-in of *Fiji/ImageJ*. Here, cells were first segmented by identifying each cell automatically in the image and by measuring Nile red and mVenus intensity profiles for each individual cell, perpendicular to the long axis. Neighboring cells or cell aggregates were filtered out. The obtained data were further processed for quantitative assessment by calculating the Jensen-Shannon Divergence (JSD) by an *Rstudio* plug-in (Rstudio, PBC, Boston, USA). JSD measures the similarity of two probability distributions. It quantifies the proximity between two distribution profiles by detecting the deviations from one distribution profile to another. The divergence was calculated by comparing the distribution of mVenus with a normal distribution of Nile red. The JSD was evaluated with scores between 0 (identical distribution) and 1 (represents maximally different distribution) ^56^.

Autoradiography samples were analyzed by using the *ImageQuant* (GE Healthcare, Munich, Germany) or *ImageJ/ Fiji* plug-in software (NIH, Bethesda, USA). All experiments were performed multiple times as independent biological replicates and technical replicates as indicated in the legends to the figures and representative gels/blots/images are shown. Mean values and SEM values were determined by using either *Excel* (Microsoft Corp, Munich, Germany) or *GraphPad Prism* (GraphPad Prism Corp. San Diego, USA). For statistical analyses, a Student unpaired two-way t-test with the Satterthwaite correction was performed (Welch-test; https://matheguru.com/). Probability values (*p*-values) are indicated in the legends to the figures. A *p*-value > 0.05 was generally considered to be not significant (n.s.).

## Supplemental Information Titles and Legends

### Legends to Supplementary Figures

**Figure S1: Fluorescence distribution profile of E. coli cells expressing different mRNAs.** (**A**) *E. coli* cells producing either MS2-Venus or MS2-Venus together with the indicated mRNAs were grown as described in the legend to Fig. 1. Subsequently, fluorescence distribution of the Nile red fluorescence (blue) and the MS2-Venus fluorescence (orange) was analyzed in individual cells and is displayed as relative fluorescence intensity. Polar areas of cells were excluded and quantification was performed along a vertical axis in mid-cell as indicated in the micrograph. The Jensen-Shannon divergence (JSD) value of the analyzed cell is indicated and JSD values close to “0” indicate almost identical distributions of Nile red and MS2-venus fluorescence. The averaged JSD values of multiple cells (n > 60-80) are shown in Fig. 1. **(B)** *In vivo* localization of *yohP* mRNA in the presence and absence of the 5‵- and 3‵- UTRs. Cells just expressing MS2-Venus were used as a control. Sample preparation and imaging was performed as described in Fig.1. **(C)** Jensen-Shannon divergence of the samples in (B). (**D**) RNA-FISH of wild-type *E. coli* cells using the TAMRA-labelled *yohP* probes as described in Fig, 1C. Note that the expression of the chromosomal *yohP* transcript was too low to be clearly localized. As negative controls, the experiment was performed with TAMRA-labelled *psbA* probes, specific for the *psbA* mRNA which encodes a photosystem II protein from the cyanobacterium *Synechocystis*sp PCC6803 and without adding probes.

**Figure S2: Nucleotide sequences and predicted secondary structures of the yohP mRNAs used in this study. (A)** *yohP* mRNA variants with different nucleotide composition were designed without changes in the predicted translation product. Nucleotides that were replaced are indicated in red. U-rich*, C-rich*, A-rich* and G-rich* refer to *yohP* variants with changes in the nucleotide composition that have only a small impact on the predicted secondary structures of the mRNA. *yohP*(Δx-x) refer to *yohP* variants with loop deletions and *yohP*(x-x subs) to variants with substituted nucleotides in the mRNA loops. **(B)** Nucleotide content and length of the different *yohP* variants shown in (A). **(C)** Predicted secondary structures of the indicated mRNAs. Structures were predicted by using the *RNAfold*webserver (http://rna.tbi.univie.ac.at/cgi-bin/RNAWebSuite/RNAfold.cgi). All variants were generated by Q5 site-Directed-Mutagenesis kit and NEBuilder HiFi Assembly Master Mix.

**Figure S3: Influence of nucleotide composition and secondary structure on membrane binding of yohP mRNAs. (A)** Jenson-Shannon divergence (JSD) plot of *yohP* mRNAs with modified nucleotide composition as shown in Fig. 2 and listed in Fig. S2. The JSD was evaluated with scores between 0 (identical distribution) and 1 (represents maximally different distribution) and is based on scoring 60-80 individual cells. Individual cells are shown in Figs. 2. **(B)** Total RNA isolated from cells expressing different mRNAs and separated by agarose gel electrophoresis. 2 µg total RNA were loaded and the gel was stained with Sybr green. **(C)** & **(D)** Northern-blot analyses of different *yohP* mRNAs. The total RNA (2 µg) shown in C was blotted onto a nylon membrane and decorated with a ^32^P-labelled oligonucleotide, complementary to the MS2-stem loop. **(E)** Quantification of three independent Northern blot experiments was performed by using *Image J* and the mean values and the SEM values are indicated. **(F)** Jenson-Shannon divergence (JSD) plot of *yohP* mRNAs with loop deletions as shown in Fig. 3 and listed in Fig. S2.

**Figure S4: The *yohP* mRNAs used for imaging are not translated and bind to liposomes of different lipid composition. (A)** *In vitro* translation of *yohP* mRNAs either containing the ribosome-binding site (+RBS) or lacking it (-RBS). The *in vitro* translation system consisted of a purified cytosolic extract, purified ribosomes and an amino acid mixture containing ^35^S-labelled methionine and cysteine. Samples were separated by SDS-PAGE and analyzed by autoradiography. YohP tends to dimerize and the YohP monomer and dimer are indicated. **(B)** The *yohP* mRNA was *in vitro* transcribed and ^32^P labeled. After purification, the mRNA was incubated with liposomes (2.5 µg/µl) of different lipid composition (PE, phosphatidyl-ethanolamine, PC, phosphatidyl-choline; CL; cardiolipin; PG, phosphatidyl-glycerol). After incubation, liposomes and the bound mRNA were pelleted by centrifugation and the membrane fraction (P) and the soluble fraction (S) were separated on a urea-PAGE-gel and analyzed by autoradiography. **(C)** Quantification of the data shown in (B). Shown are the mean values of at least three experiments and SEM in indicated by error bars. Statistical analyses were performed with the Satterthwaite corrected unpaired two-sided Student t-test, using the values after incubation without INVs as reference. (*) refers to *p*-values ≤0.05; (**) to *p*-values ≤0.01, and (***) to *p*-values ≤0.001. **(D)** Immune detection of SecY in different INVs. SecYEG-OE refers to INVs from a SecYEG-overproducing strain and SecY(ΔC4-6)EG to INVs from an *E. coli* strain that overproduces the truncated SecYEG complex. Antibodies against YidC were used as loading control. **(E)** Cartoon showing the TM-MS2-venus construct (left panel) and a microscopic image showing *E. coli* cells producing TM2-MS2-venus (right panel). The cartoon was generated using Biorender (https://biorender.com/).

**Figure S5: Localization of different SecY-variants in the *E. coli* membrane**. **(A)** Different variants of the SecYEG translocon with a C-terminally YFP-tagged SecY were produced in *E. coli* and membrane localization was analyzed by fluorescence microscopy. ΔC4, ΔC5, Δ6 and ΔC4-6 refer to SecY deletions that lack either the cytosolic loop C4, C5, C6 or all three loops, respectively. The scale bar refers to 2 µM. **(B**) Immune detection of YidC in wild type INVs and INVs from a YidC overproducing strain. The small size shift is due to the presence of a His-tag in the plasmid-encoded YidC. **(C)** *In vitro* translation of *yohP* variants that either contain or lack the MS2 stem loop. *In vitro* translation was performed as described in Fig. S4 in the presence of wild type INV and one half of the reaction was treated with proteinase K for detecting membrane inserted YohP.

**Figure S6: mRNA targeting is translation independent. (A-D)** For validating that mRNA targeting occurs independently of translation, mRNA targeting was followed in the absence or presence of antibiotics. Kasugamycin (0.1 mg/ml) or Puromycin (0.2 mg/ml) were present during the last 10 min of MS2-Venus induction. Microscopy was performed as described in the legend to Fig. 1. **(E)** JSD-plot of 60-80 individual cells, treated as in (A-D).

**Figure S7: Validation of the experimental strategy for determining the role of SRP in membrane protein insertion. (A)** Urea treatment removes large amounts of FtsY and SRP (Ffh) from INVs. Urea-treated INVs (U-INVs) were generated by treating INVs with 6 M urea; after incubation and centrifugation, U-INV (2.5 µg) were separated by SDS-PAGE and after western-blotting decorated with antibodies against Ffh, the protein component of the bacterial SRP, and FtsY, the bacterial SRP receptor. Untreated INVs served as control. **(B)** Pulse-chase experiment for monitoring the fractionation procedure. The cytosolic protein YchF served as a control to ensure that the procedure described in the legend to Fig.6 allows to separate soluble from inner membrane proteins. Cells were grown on minimal media and YchF production was induced by the addition of IPTG. ^35^S-labelled methionine/cysteine mix was then added, followed by 5 min incubation. Cells were chased for 1, 5 and 10 min with an excess of non-radioactive methionine and cysteine. Cells were then lysed by sonication as described in the legend to Fig. 6 and separated by ultracentrifugation in the soluble fraction and the membrane fraction. Both fractions were then separated on SDS-PAGE and the gel was analyzed by autoradiography. The position of 40 kDa YchF is indicated. **(C)** Norvaline inhibits global translation in *E. coli*. Cells were grown on minimal media and treated for 90 min with norvaline when indicated. ^35^S-labelled methionine/cysteine mix was then added, followed by 5 min incubation. Cells were chased for 1, 5 and 10 min with an excess of non-radioactive methionine and cysteine. After separation on SDS-PAGE, the gel was analyzed by autoradiography. YohP is indicated.

**Table S1: *E. coli* strains used in this study**

**Table S2: Plasmids used in this study**

Plasmids used for *in vivo* mRNA imaging lacked the Shine-Dalgarno sequence (SD).

**Table S3: Oligonucleotides used in this study**

**Table S4: *yohP* probes used in this study**

## References

1. Driessen, A.J., Manting, E.H., and van der Does, C. (2001). The structural basis of protein targeting and translocation in bacteria. Nature structural biology 8, 492–498. 10.1038/88549.

2. Muller, M., Koch, H.G., Beck, K., and Schafer, U. (2001). Protein traffic in bacteria: multiple routes from the ribosome to and across the membrane. Progress in nucleic acid research and molecular biology 66, 107–157.

3. Rapoport, T.A., Li, L., and Park, E. (2017). Structural and Mechanistic Insights into Protein Translocation. Annual review of cell and developmental biology 33, 369–390. 10.1146/annurev-cellbio-100616-060439.

4. Zimmermann, R., Eyrisch, S., Ahmad, M., and Helms, V. (2011). Protein translocation across the ER membrane. Biochimica et biophysica acta 1808, 912–924.

5. Oswald, J., Njenga, R., Natriashvili, A., Sarmah, P., and Koch, H.G. (2021). The Dynamic SecYEG Translocon. Front Mol Biosci 8, 664241. 10.3389/fmolb.2021.664241.

6. Denks, K., Vogt, A., Sachelaru, I., Petriman, N.A., Kudva, R., and Koch, H.G. (2014). The Sec translocon mediated protein transport in prokaryotes and eukaryotes. Molecular membrane biology 31, 58–84.

7. Osborne, A.R., Rapoport, T.A., and van den Berg, B. (2005). Protein translocation by the Sec61/SecY channel. Annual review of cell and developmental biology 21, 529–550. 10.1146/annurev.cellbio.21.012704.133214.

8. Vögtle, F.N., Koch, H.G., and Meisinger, C. (2022). A common evolutionary origin reveals fundamental principles of protein insertases. PLoS biology 20, e3001558. 10.1371/journal.pbio.3001558.

9. McDowell, M.A., Heimes, M., and Sinning, I. (2021). Structural and molecular mechanisms for membrane protein biogenesis by the Oxa1 superfamily. Nature structural & molecular biology 28, 234–239. 10.1038/s41594-021-00567-9.

10. Dalbey, R., Koch, H.G., and Kuhn, A. (2017). Targeting and Insertion of Membrane Proteins. EcoSalPlus 7, 1–28.

11. Welte, T., Kudva, R., Kuhn, P., Sturm, L., Braig, D., Muller, M., Warscheid, B., Drepper, F., and Koch, H.G. (2012). Promiscuous targeting of polytopic membrane proteins to SecYEG or YidC by the Escherichia coli signal recognition particle. Molecular biology of the cell 23, 464–479. 10.1091/mbc.E11-07-0590.

12. Steinberg, R., Knupffer, L., Origi, A., Asti, R., and Koch, H.G. (2018). Co-translational protein targeting in bacteria. FEMS microbiology letters 365, 1–15. 10.1093/femsle/fny095.

13. Akopian, D., Shen, K., Zhang, X., and Shan, S.O. (2013). Signal recognition particle: an essential protein-targeting machine. Annual review of biochemistry 82, 693–721. 10.1146/annurev-biochem-072711-164732.

14. Herskovits, A.A., Bochkareva, E.S., and Bibi, E. (2000). New prospects in studying the bacterial signal recognition particle pathway. Molecular microbiology 38, 927–939.

15. Rodnina, M.V., and Wintermeyer, W. (2016). Protein Elongation, Co-translational Folding and Targeting. Journal of molecular biology 428, 2165–2185. 10.1016/j.jmb.2016.03.022.

16. Denks, K., Sliwinski, N., Erichsen, V., Borodkina, B., Origi, A., and Koch, H.G. (2017). The signal recognition particle contacts uL23 and scans substrate translation inside the ribosomal tunnel. Nature Microbiology 2, 16265. 10.1028/nmicrobiol.2016.265.

17. Schaffitzel, C., Oswald, M., Berger, I., Ishikawa, T., Abrahams, J.P., Koerten, H.K., Koning, R.I., and Ban, N. (2006). Structure of the E. coli signal recognition particle bound to a translating ribosome. Nature 444, 503–506. 10.1038/nature05182.

18. Halic, M., Becker, T., Pool, M.R., Spahn, C.M., Grassucci, R.A., Frank, J., and Beckmann, R. (2004). Structure of the signal recognition particle interacting with the elongation-arrested ribosome. Nature 427, 808–814. 10.1038/nature02342.

19. Gu, S.Q., Peske, F., Wieden, H.J., Rodnina, M.V., and Wintermeyer, W. (2003). The signal recognition particle binds to protein L23 at the peptide exit of the Escherichia coli ribosome. RNA (New York, N.Y.) 9, 566–573.

20. Jomaa, A., Fu, Y.H., Boehringer, D., Leibundgut, M., Shan, S.O., and Ban, N. (2017). Structure of the quaternary complex between SRP, SR, and translocon bound to the translating ribosome. Nature communications 8, 15470.

21. Luirink, J., ten Hagen-Jongman, C.M., van der Weijden, C.C., Oudega, B., High, S., Dobberstein, B., and Kusters, R. (1994). An alternative protein targeting pathway in Escherichia coli: studies on the role of FtsY. The EMBO journal 13, 2289–2296.

22. Peluso, P., Shan, S.O., Nock, S., Herschlag, D., and Walter, P. (2001). Role of SRP RNA in the GTPase cycles of Ffh and FtsY. Biochemistry 40, 15224–15233.

23. Braig, D., Bar, C., Thumfart, J.O., and Koch, H.G. (2009). Two cooperating helices constitute the lipid-binding domain of the bacterial SRP receptor. Journal of molecular biology 390, 401–413. 10.1016/j.jmb.2009.04.061.

24. Erez, E., Stjepanovic, G., Zelazny, A.M., Brugger, B., Sinning, I., and Bibi, E. (2010). Genetic evidence for functional interaction of the Escherichia coli signal recognition particle receptor with acidic lipids in vivo. The Journal of biological chemistry 285, 40508–40514. 10.1074/jbc.M110.140921.

25. de Leeuw, E., te Kaat, K., Moser, C., Menestrina, G., Demel, R., de Kruijff, B., Oudega, B., Luirink, J., and Sinning, I. (2000). Anionic phospholipids are involved in membrane association of FtsY and stimulate its GTPase activity. The EMBO journal 19, 531–541. 10.1093/emboj/19.4.531.

26. Weiche, B., Burk, J., Angelini, S., Schiltz, E., Thumfart, J.O., and Koch, H.G. (2008). A cleavable N-terminal membrane anchor is involved in membrane binding of the Escherichia coli SRP receptor. Journal of molecular biology 377, 761–773. 10.1016/j.jmb.2008.01.040.

27. Angelini, S., Boy, D., Schiltz, E., and Koch, H.G. (2006). Membrane binding of the bacterial signal recognition particle receptor involves two distinct binding sites. The Journal of cell biology 174, 715–724. 10.1083/jcb.200606093.

28. Angelini, S., Deitermann, S., and Koch, H.G. (2005). FtsY, the bacterial signal-recognition particle receptor, interacts functionally and physically with the SecYEG translocon. EMBO reports 6, 476–481. 10.1038/sj.embor.7400385.

29. Kuhn, P., Weiche, B., Sturm, L., Sommer, E., Drepper, F., Warscheid, B., Sourjik, V., and Koch, H.G. (2011). The bacterial SRP receptor, SecA and the ribosome use overlapping binding sites on the SecY translocon. Traffic (Copenhagen, Denmark) 12, 563–578. 10.1111/j.1600-0854.2011.01167.x.

30. Petriman, N.A., Jauss, B., Hufnagel, A., Franz, L., Sachelaru, I., Drepper, F., Warscheid, B., and Koch, H.G. (2018). The interaction network of the YidC insertase with the SecYEG translocon, SRP and the SRP receptor FtsY. Scientific reports 8, 578. 10.1038/s41598-017-19019-w.

31. Draycheva, A., Bornemann, T., Ryazanov, S., Lakomek, N.A., and Wintermeyer, W. (2016). The bacterial SRP receptor, FtsY, is activated on binding to the translocon. Molecular microbiology 102, 152–167. 10.1111/mmi.13452.

32. Kuhn, P., Draycheva, A., Vogt, A., Petriman, N.A., Sturm, L., Drepper, F., Warscheid, B., Wintermeyer, W., and Koch, H.G. (2015). Ribosome binding induces repositioning of the signal recognition particle receptor on the translocon. The Journal of cell biology 211, 91–104. 10.1111/mmi.13321.

33. Halic, M., Gartmann, M., Schlenker, O., Mielke, T., Pool, M.R., Sinning, I., and Beckmann, R. (2006). Signal recognition particle receptor exposes the ribosomal translocon binding site. Science (New York, N.Y.) 312, 745–747. 10.1126/science.1124864.

34. White, S.H., and von Heijne, G. (2004). The machinery of membrane protein assembly. Current opinion in structural biology 14, 397–404. 10.1016/j.sbi.2004.07.003.

35. Cymer, F., von Heijne, G., and White, S.H. (2015). Mechanisms of integral membrane protein insertion and folding. Journal of molecular biology 427, 999–1022. 10.1016/j.jmb.2014.09.014.

36. Bradshaw, N., Neher, S.B., Booth, D.S., and Walter, P. (2009). Signal sequences activate the catalytic switch of SRP RNA. Science (New York, N.Y.) 323, 127–130. 10.1126/science.1165971.

37. Corey, R.A., Allen, W.J., Komar, J., Masiulis, S., Menzies, S., Robson, A., and Collinson, I. (2016). Unlocking the Bacterial SecY Translocon. Structure (London, England : 1993) *24*, 518–527. 10.1016/j.str.2016.02.001.

38. Goder, V., and Spiess, M. (2003). Molecular mechanism of signal sequence orientation in the endoplasmic reticulum. The EMBO journal 22, 3645–3653. 10.1093/emboj/cdg361.

39. Halic, M., Blau, M., Becker, T., Mielke, T., Pool, M.R., Wild, K., Sinning, I., and Beckmann, R. (2006). Following the signal sequence from ribosomal tunnel exit to signal recognition particle. Nature 444, 507–511. 10.1038/nature05326.

40. Hegde, R.S., and Bernstein, H.D. (2006). The surprising complexity of signal sequences. Trends in biochemical sciences 31, 563–571. 10.1016/j.tibs.2006.08.004.

41. Schibich, D., Gloge, F., Pohner, I., Bjorkholm, P., Wade, R.C., von Heijne, G., Bukau, B., and Kramer, G. (2016). Global profiling of SRP interaction with nascent polypeptides. Nature 536, 219–223. 10.1038/nature19070.

42. Nevo-Dinur, K., Nussbaum-Shochat, A., Ben-Yehuda, S., and Amster-Choder, O. (2011). Translation-independent localization of mRNA in E. coli. Science (New York, N.Y.) 331, 1081–1084.

43. Kannaiah, S., Livny, J., and Amster-Choder, O. (2019). Spatiotemporal Organization of the E. coli Transcriptome: Translation Independence and Engagement in Regulation. Molecular cell 76, 574–589 e577. 10.1016/j.molcel.2019.08.013.

44. Fei, J., and Sharma, C.M. (2018). RNA Localization in Bacteria. Microbiology spectrum 6. 101128/microbiolspec.RWR-0024-2018.

45. van Gijtenbeek, L.A., Robinson, A., van Oijen, A.M., Poolman, B., and Kok, J. (2016). On the Spatial Organization of mRNA, Plasmids, and Ribosomes in a Bacterial Host Overexpressing Membrane Proteins. PLoS genetics 12, e1006523. 10.1371/journal.pgen.1006523.

46. Moffitt, J.R., Pandey, S., Boettiger, A.N., Wang, S., and Zhuang, X. (2016). Spatial organization shapes the turnover of a bacterial transcriptome. Elife 5. 10.7554/eLife.13065.

47. Prilusky, J., and Bibi, E. (2009). Studying membrane proteins through the eyes of the genetic code revealed a strong uracil bias in their coding mRNAs. Proceedings of the National Academy of Sciences of the United States of America 106, 6662–6666.

48. Kannaiah, S., and Amster-Choder, O. (2014). Protein targeting via mRNA in bacteria. Biochimica et biophysica acta 1843, 1457–1465. 10.1016/j.bbamcr.2013.11.004.

49. Benhalevy, D., Bochkareva, E.S., Biran, I., and Bibi, E. (2015). Model Uracil-Rich RNAs and Membrane Protein mRNAs Interact Specifically with Cold Shock Proteins in Escherichia coli. PloS one 10, e0134413.

50. Mahbub, M., Hemm, L., Yang, Y., Kaur, R., Carmen, H., Engl, C., Huokko, T., Riediger, M., Watanabe, S., Liu, L.N., et al. (2020). mRNA localization, reaction centre biogenesis and thylakoid membrane targeting in cyanobacteria. Nat Plants 6, 1179–1191. 10.1038/s41477-020-00764-2.

51. Benhalevy, D., Biran, I., Bochkareva, E.S., Sorek, R., and Bibi, E. (2017). Evidence for a cytoplasmic pool of ribosome-free mRNAs encoding inner membrane proteins in Escherichia coli. PloS one 12, e0183862.

52. Kannaiah, S., and Amster-Choder, O. (2016). Methods for studying RNA localization in bacteria. Methods (San Diego, Calif.) 98, 99–103. 10.1016/j.ymeth.2015.12.010.

53. Steinberg, R., Origi, A., Natriashvili, A., Sarmah, P., Licheva, M., Walker, P.M., Kraft, C., High, S., Luirink, J., Shi, W.Q., et al. (2020). Posttranslational insertion of small membrane proteins by the bacterial signal recognition particle. PLoS biology 18, e3000874. 10.1371/journal.pbio.3000874.

54. Valax, P., and Georgiou, G. (1993). Molecular characterization of beta-lactamase inclusion bodies produced in Escherichia coli. 1. Composition. Biotechnol. Prog. 9, 539–547.

55. Koch, H.G., and Muller, M. (2000). Dissecting the translocase and integrase functions of the Escherichia coli SecYEG translocon. The Journal of cell biology 150, 689–694.

56. Nielsen, F. (2020). On a Generalization of the Jensen-Shannon Divergence and the Jensen-Shannon Centroid. Entropy (Basel, Switzerland) 22. 10.3390/e22020221.

57. Cohen-Zontag, O., Baez, C., Lim, L.Q.J., Olender, T., Schirman, D., Dahary, D., Pilpel, Y., and Gerst, J.E. (2019). A secretion-enhancing cis regulatory targeting element (SECReTE) involved in mRNA localization and protein synthesis. PLoS genetics 15, e1008248. 10.1371/journal.pgen.1008248.

58. Pannwitt, S., Slama, K., Depoix, F., Helm, M., and Schneider, D. (2019). Against Expectations: Unassisted RNA Adsorption onto Negatively Charged Lipid Bilayers. Langmuir : the ACS journal of surfaces and colloids 35, 14704–14711. 10.1021/acs.langmuir.9b02830.

59. Dabkowska, A.P., Michanek, A., Jaeger, L., Chworos, A., Nylander, T., and Sparr, E. (2017). Supported Fluid Lipid Bilayer as a Scaffold to Direct Assembly of RNA Nanostructures. Methods in molecular biology (Clifton, N.J.) 1632, 107–122. 10.1007/978-1-4939-7138-1_7.

60. Michanek, A., Yanez, M., Wacklin, H., Hughes, A., Nylander, T., and Sparr, E. (2012). RNA and DNA association to zwitterionic and charged monolayers at the air-liquid interface. Langmuir : the ACS journal of surfaces and colloids 28, 9621–9633. 10.1021/la204431q.

61. Czerniak, T., and Saenz, J.P. (2022). Lipid membranes modulate the activity of RNA through sequence-dependent interactions. Proceedings of the National Academy of Sciences of the United States of America 119. 10.1073/pnas.2119235119.

62. Peabody, D.S. (1993). The RNA binding site of bacteriophage MS2 coat protein. The EMBO journal 12, 595–600.

63. Pyhtila, B., Zheng, T., Lager, P.J., Keene, J.D., Reedy, M.C., and Nicchitta, C.V. (2008). Signal sequence- and translation-independent mRNA localization to the endoplasmic reticulum. RNA (New York, N.Y.) 14, 445–453. 10.1261/rna.721108.

64. Jagannathan, S., Hsu, J.C., Reid, D.W., Chen, Q., Thompson, W.J., Moseley, A.M., and Nicchitta, C.V. (2014). Multifunctional roles for the protein translocation machinery in RNA anchoring to the endoplasmic reticulum. The Journal of biological chemistry 289, 25907–25924. 10.1074/jbc.M114.580688.

65. Bhadra, P., Schorr, S., Lerner, M., Nguyen, D., Dudek, J., Förster, F., Helms, V., Lang, S., and Zimmermann, R. (2021). Quantitative Proteomics and Differential Protein Abundance Analysis after Depletion of Putative mRNA Receptors in the ER Membrane of Human Cells Identifies Novel Aspects of mRNA Targeting to the ER. Molecules (Basel, Switzerland) 26. 10.3390/molecules26123591.

66. Carpousis, A.J. (2007). The RNA degradosome of Escherichia coli: an mRNA-degrading machine assembled on RNase E. Annual review of microbiology 61, 71–87. 10.1146/annurev.micro.61.080706.093440.

67. Mori, H., and Ito, K. (2006). Different modes of SecY-SecA interactions revealed by site-directed in vivo photo-cross-linking. Proceedings of the National Academy of Sciences of the United States of America 103, 16159–16164. 10.1073/pnas.0606390103.

68. Das, S., and Oliver, D.B. (2011). Mapping of the SecA.SecY and SecA.SecG interfaces by site-directed in vivo photocross-linking. The Journal of biological chemistry 286, 12371–12380. 10.1074/jbc.M110.182931.

69. Karamanou, S., Bariami, V., Papanikou, E., Kalodimos, C.G., and Economou, A. (2008). Assembly of the translocase motor onto the preprotein-conducting channel. Molecular microbiology 70, 311–322. 10.1111/j.1365-2958.2008.06402.x.

70. Fessl, T., Watkins, D., Oatley, P., Allen, W.J., Corey, R.A., Horne, J., Baldwin, S.A., Radford, S.E., Collinson, I., and Tuma, R. (2018). Dynamic action of the Sec machinery during initiation, protein translocation and termination. Elife 7.

71. Kusters, I., van den Bogaart, G., Kedrov, A., Krasnikov, V., Fulyani, F., Poolman, B., and Driessen, A.J. (2011). Quaternary structure of SecA in solution and bound to SecYEG probed at the single molecule level. Structure (London, England : 1993) 19, 430–439. 10.1016/j.str.2010.12.016.

72. Zimmer, J., Nam, Y., and Rapoport, T.A. (2008). Structure of a complex of the ATPase SecA and the protein-translocation channel. Nature 455, 936–943. 10.1038/nature07335.

73. Prinz, A., Behrens, C., Rapoport, T.A., Hartmann, E., and Kalies, K.U. (2000). Evolutionarily conserved binding of ribosomes to the translocation channel via the large ribosomal RNA. The EMBO journal 19, 1900–1906. 10.1093/emboj/19.8.1900.

74. Cheng, Z., Jiang, Y., Mandon, E.C., and Gilmore, R. (2005). Identification of cytoplasmic residues of Sec61p involved in ribosome binding and cotranslational translocation. The Journal of cell biology 168, 67–77. 10.1083/jcb.200408188.

75. Jomaa, A., Boehringer, D., Leibundgut, M., and Ban, N. (2016). Structure of the E. coli translating ribosome with SRP and its receptor and with the translocon. Nat. Comm. 7, 10471.

76. Frauenfeld, J., Gumbart, J., Sluis, E.O., Funes, S., Gartmann, M., Beatrix, B., Mielke, T., Berninghausen, O., Becker, T., Schulten, K., and Beckmann, R. (2011). Cryo-EM structure of the ribosome-SecYE complex in the membrane environment. Nature structural & molecular biology 18, 614–621. 10.1038/nsmb.2026.

77. Bischoff, L., Wickles, S., Berninghaus, O., van der Sluis, E.O., and Beckmann, R. (2014). Visualization of a polytopic membrane protein during SecY-mediated membrane insertion. Nature communications 5, 4103.

78. Bürk, J., Weiche, B., Wenk, M., Boy, D., Nestel, S., Heimrich, B., and Koch, H.G. (2009). Depletion of the signal recognition particle receptor inactivates ribosomes in Escherichia coli. J. Bacteriol. 191, 7017–7026.

79. Baars, L., Wagner, S., Wickstrom, D., Klepsch, M., Ytterberg, A.J., van Wijk, K.J., and de Gier, J.W. (2008). Effects of SecE depletion on the inner and outer membrane proteomes of Escherichia coli. Journal of bacteriology 190, 3505–3525.

80. Wickstrom, D., Wagner, S., Simonsson, P., Pop, O., Baars, L., Ytterberg, A.J., van Wijk, K.J., Luirink, J., and de Gier, J.W. (2011). Characterization of the consequences of YidC depletion on the inner membrane proteome of E. coli using 2D blue native/SDS-PAGE. Journal of molecular biology 409, 124–135. 10.1016/j.jmb.2011.03.068.

81. Wang, P., Kuhn, A., and Dalbey, R.E. (2010). Global change of gene expression and cell physiology in YidC-depleted Escherichia coli. Journal of bacteriology 192, 2193–2209. 10.1128/jb.00484-09.

82. Price, C.E., Otto, A., Fusetti, F., Becher, D., Hecker, M., and Driessen, A.J. (2010). Differential effect of YidC depletion on the membrane proteome of Escherichia coli under aerobic and anaerobic growth conditions. Proteomics 10, 3235–3247. 10.1002/pmic.201000284.

83. Kuhn, P., Kudva, R., Welte, T., Sturm, L., and Koch, H.G. (2013). Targeting and Integration of bacterial membrane proteins. Bacterial Membranes: Structural and Molecular Biology (Remaut, H. & Fronzes, R., Eds), 303–343.

84. Geng, Y., Kedrov, A., Caumanns, J.J., Crevenna, A.H., Lamb, D.C., Beckmann, R., and Driessen, A.J. (2015). Role of the Cytosolic Loop C2 and the C Terminus of YidC in Ribosome Binding and Insertion Activity. The Journal of biological chemistry 290, 17250–17261. 10.1074/jbc.M115.650309.

85. Wickles, S., Singharoy, A., Andreani, J., Seemayer, S., Bischoff, L., Berninghausen, O., Soeding, J., Schulten, K., van der Sluis, E.O., and Beckmann, R. (2014). A structural model of the active ribosome-bound membrane protein insertase YidC. Elife 3, e03035. 10.7554/eLife.03035.

86. Kohler, R., Boehringer, D., Greber, B., Bingel-Erlenmeyer, R., Collinson, I., Schaffitzel, C., and Ban, N. (2009). YidC and Oxa1 form dimeric insertion pores on the translating ribosome. Molecular cell 34, 344–353. 10.1016/j.molcel.2009.04.019.

87. El-Sharoud, W.M., and Graumann, P.L. (2007). Cold shock proteins aid coupling of transcription and translation in bacteria. Science Progress, 15–27.

88. Koch, H.G., Hengelage, T., Neumann-Haefelin, C., MacFarlane, J., Hoffschulte, H.K., Schimz, K.L., Mechler, B., and Muller, M. (1999). In vitro studies with purified components reveal signal recognition particle (SRP) and SecA/SecB as constituents of two independent protein-targeting pathways of Escherichia coli. Molecular biology of the cell 10, 2163–2173.

89. Bornemann, T., Jockel, J., Rodnina, M.V., and Wintermeyer, W. (2008). Signal sequence-independent membrane targeting of ribosomes containing short nascent peptides within the exit tunnel. Nature structural & molecular biology 15, 494–499. 10.1038/nsmb.1402.

90. Koch, H.G., Moser, M., Schimz, K.L., and Muller, M. (2002). The integration of YidC into the cytoplasmic membrane of Escherichia coli requires the signal recognition particle, SecA and SecYEG. The Journal of biological chemistry 277, 5715–5718. 10.1074/jbc.C100683200.

91. Czech, L., Mais, C.N., Kratzat, H., Sarmah, P., Giammarinaro, P., Freibert, S.A., Esser, H.F., Musial, J., Berninghausen, O., Steinchen, W., et al. (2022). Inhibition of SRP-dependent protein secretion by the bacterial alarmone (p)ppGpp. Nature communications 13, 1069. 10.1038/s41467-022-28675-0.

92. Kudva, R., Denks, K., Kuhn, P., Vogt, A., Muller, M., and Koch, H.G. (2013). Protein translocation across the inner membrane of Gram-negative bacteria: the Sec and Tat dependent protein transport pathways. Research in microbiology 164, 505–534. 10.1016/j.resmic.2013.03.016.

93. Cvetesic, N., Palencia, A., Halasz, I., Cusack, S., and Gruic-Sovulj, I. (2014). The physiological target for LeuRS translational quality control is norvaline. The EMBO journal 33, 1639–1653.

94. Eymann, C., Mittenhuber, G., and Hecker, M. (2001). The stringent response, sigmaH-dependent gene expression and sporulation in Bacillus subtilis. Molecular & general genetics : MGG 264, 913–923. 10.1007/s004380000381.

95. Schäfer, H., Beckert, B., Frese, C.K., Steinchen, W., Nuss, A.M., Beckstette, M., Hantke, I., Driller, K., Sudzinová, P., Krásný, L., et al. (2020). The alarmones (p)ppGpp are part of the heat shock response of Bacillus subtilis. PLoS genetics 16, e1008275.

96. Steinchen, W., Zegarra, V., and Bange, G. (2020). (p)ppGpp: Magic Modulators of Bacterial Physiology and Metabolism. Front Microbiol 11, 2072. 10.3389/fmicb.2020.02072.

97. Tabib-Salazar, A., Liu, B., Barker, D., Burchell, L., Qimron, U., Matthews, S.J., and Wigneshweraraj, S. (2018). T7 phage factor required for managing RpoS in Escherichia coli. Proceedings of the National Academy of Sciences of the United States of America 115, E5353–E5362.

98. Mutka, S.C., and Walter, P. (2001). Multifaceted physiological response allows yeast to adapt to the loss of the signal recognition particle-dependent protein-targeting pathway. Molecular biology of the cell 12, 577–588. 10.1091/mbc.12.3.577.

99. Bernstein, H.D., and Hyndman, J.B. (2001). Physiological basis for conservation of the signal recognition particle targeting pathway in Escherichia coli. Journal of bacteriology 183, 2187–2197. 10.1128/jb.183.7.2187-2197.2001.

100. Zhao, L., Fu, G., Cui, Y., Xu, Z., Cai, T., and Zhang, D. (2021). Compensating Complete Loss of Signal Recognition Particle During Co-translational Protein Targeting by the Translation Speed and Accuracy. Front Microbiol 12, 690286. 10.3389/fmicb.2021.690286.

101. Landwehr, V., Milanov, M., Angebauer, L., Hong, J., Jüngert, G., Hiersemenzel, A., Siebler, A., Schmit, F., Öztürk, Y., Dannenmaier, S., et al. (2021). The Universally Conserved ATPase YchF Regulates Translation of Leaderless mRNA in Response to Stress Conditions. Front Mol Biosci 8, 643696. 10.3389/fmolb.2021.643696.

102. Beck, H.J., and Moll, I. (2018). Leaderless mRNAs in the Spotlight: Ancient but Not Outdated! Microbiology spectrum 6. 10.1128/microbiolspec.RWR-0016-2017.

103. Hasona, A., Crowley, P.J., Levesque, C.M., Mair, R.W., Cvitkovitch, D.G., Bleiweis, A.S., and Brady, L.J. (2005). Streptococcal viability and diminished stress tolerance in mutants lacking the signal recognition particle pathway or YidC2. Proceedings of the National Academy of Sciences of the United States of America 102, 17466–17471. 10.1073/pnas.0508778102.

104. Palmer, S.R., Crowley, P.J., Oli, M.W., Ruelf, M.A., Michalek, S.M., and Brady, L.J. (2012). YidC1 and YidC2 are functionally distinct proteins involved in protein secretion, biofilm formation and cariogenicity of Streptococcus mutans. Microbiology (Reading, England) 158, 1702–1712. 10.1099/mic.0.059139-0.

105. Fouts, D.E., Matthias, M.A., Adhikarla, H., Adler, B., Amorim-Santos, L., Berg, D.E., Bulach, D., Buschiazzo, A., Chang, Y.F., Galloway, R.L., et al. (2016). What Makes a Bacterial Species Pathogenic?:Comparative Genomic Analysis of the Genus Leptospira. PLoS neglected tropical diseases 10, e0004403.

106. Sauerbrei, B., Arends, J., Schünemann, D., and Narberhaus, F. (2020). Lon Protease Removes Excess Signal Recognition Particle Protein in Escherichia coli. Journal of bacteriology 202. 10.1128/jb.00161-20.

107. Steinberg, R., and Koch, H.G. (2021). The largely unexplored biology of small proteins in pro- and eukaryotes. The FEBS journal. 10.1111/febs.15845.

108. Fontaine, F., Fuchs, R.T., and Storz, G. (2011). Membrane localization of small proteins in Escherichia coli. The Journal of biological chemistry 286, 32464–32474.

109. Hemm, M.R., Paul, B.J., Miranda-Rios, J., Zhang, A., Soltanzad, N., and Storz, G. (2010). Small stress response proteins in Escherichia coli: proteins missed by classical proteomic studies. Journal of bacteriology 192, 46–58. 10.1093/nar/gkq054.

110. Hobbs, E.C., Astarita, J.L., and Storz, G. (2010). Small RNAs and small proteins involved in resistance to cell envelope stress and acid shock in Escherichia coli: analysis of a bar-coded mutant collection. Journal of bacteriology 192, 59–67. 10.1128/jb.00873-09.

111. Behrens, C., Hartmann, E., and Kalies, K.U. (2013). Single rRNA helices bind independently to the protein-conducting channel SecYEG. Traffic (Copenhagen, Denmark) 14, 274–281. 10.1111/tra.12033.

112. Kedrov, A., Wickles, S., Crevenna, A.H., van der Sluis, E.O., Buschauer, R., Berninghausen, O., Lamb, D.C., and Beckmann, R. (2016). Structural Dynamics of the YidC:Ribosome Complex during Membrane Protein Biogenesis. Cell reports 17, 2943–2954. 10.3389/fnana.2017.00117.

113. Seitl, I., Wickles, S., Beckmann, R., Kuhn, A., and Kiefer, D. (2014). The C-terminal regions of YidC from Rhodopirellula baltica and Oceanicaulis alexandrii bind to ribosomes and partially substitute for SRP receptor function in Escherichia coli. Molecular microbiology 91, 408–421. 10.1111/mmi.12465.

114. Miller, O.L., Jr., Hamkalo, B.A., and Thomas, C.A., Jr. (1970). Visualization of bacterial genes in action. Science (New York, N.Y.) 169, 392–395. 10.1126/science.169.3943.392.

115. Kohler, R., Mooney, R.A., Mills, D.J., Landick, R., and Cramer, P. (2017). Architecture of a transcribing-translating expressome. Science (New York, N.Y.) 356, 194–197.

116. Irastortza-Olaziregi, M., and Amster-Choder, O. (2020). Coupled Transcription-Translation in Prokaryotes: An Old Couple With New Surprises. Front Microbiol 11, 624830. 10.3389/fmicb.2020.624830.

117. Anderson, D.M., and Schneewind, O. (1997). A mRNA signal for the type III secretion of Yop proteins by Yersinia enterocolitica. Science (New York, N.Y.) 278, 1140–1143. 10.1126/science.278.5340.1140.

118. Skinner, S.O., Sepúlveda, L.A., Xu, H., and Golding, I. (2013). Measuring mRNA copy number in individual Escherichia coli cells using single-molecule fluorescent in situ hybridization. Nature protocols 8, 1100–1113.

119. Pinto, F.L., Thapper, A., Sontheim, W., and Lindblad, P. (2009). Analysis of current and alternative phenol based RNA extraction methodologies for cyanobacteria. BMC Mol Biol 10, 79.

120. Hoffschulte, H.K., Drees, B., and Muller, M. (1994). Identification of a soluble SecA/SecB complex by means of a subfractionated cell-free export system. The Journal of biological chemistry 269, 12833–12839.

121. Braig, D., Mircheva, M., Sachelaru, I., van der Sluis, E.O., Sturm, L., Beckmann, R., and Koch, H.G. (2011). Signal sequence-independent SRP-SR complex formation at the membrane suggests an alternative targeting pathway within the SRP cycle. Molecular biology of the cell 22, 2309–2323.

122. Schägger, H. (2006). Tricine-SDS-PAGE. Nature protocols 1, 16–22. 10.1038/nprot.2006.4.

123. Boy, D., and Koch, H.G. (2009). Visualization of distinct entities of the SecYEG translocon during translocation and integration of bacterial proteins. Molecular biology of the cell 20, 1804–1815. 10.1091/mbc.E08-08-0886.

