## Supplemental information for "mRNA targeting eliminates the need for the signal recognition particle during membrane protein insertion in bacteria"

**Figure S7: Validation of the experimental strategy for determining the role of SRP in membrane protein insertion. (A)** Urea treatment removes large amounts of FtsY and SRP (Ffh) from INVs. Urea-treated INVs (U-INVs) were generated by treating INVs with 6 M urea; after incubation and centrifugation, U-INV (2.5 µg) were separated by SDS-PAGE and after western-blotting decorated with antibodies against Ffh, the protein component of the bacterial SRP, and FtsY, the bacterial SRP receptor. Untreated INVs served as control. **(B)** Pulse-chase experiment for monitoring the fractionation procedure. The cytosolic protein YchF served as a control to ensure that the procedure described in the legend to Fig.6 allows to separate soluble from inner membrane proteins. Cells were grown on minimal media and YchF production was induced by the addition of IPTG. $^{35}$S-labelled methionine/cysteine mix was then added, followed by 5 min incubation. Cells were chased for 1, 5 and 10 min with an excess of non-radioactive methionine and cysteine. Cells were then lysed by sonication as described in the legend to Fig. 6 and separated by ultracentrifugation in the soluble fraction and the membrane fraction. Both fractions were then separated on SDS-PAGE and the gel was analyzed by autoradiography. The position of 40 kDa YchF is indicated. **(C)** Norvaline inhibits global translation in *E. coli*. Cells were grown on minimal media and treated for 90 min with norvaline when indicated. $^{35}$S-labelled methionine/cysteine mix was then added, followed by 5 min incubation. Cells were chased for 1, 5 and 10 min with an excess of non-radioactive methionine and cysteine. After separation on SDS-PAGE, the gel was analyzed by autoradiography. YohP is indicated.

**Table S1: *E. coli* strains used in this study**

| Name | Description | Application | Reference |
|---|---|---|---|
| DH5α | *supE44 ΔlacU169 (Φ80lacZΔM15)hsdR17 recA1 endA1 gyrA96 thi-1 relA1* | Plasmid isolation and storage | [1] |
| BL21 | *E. coli B F- dcm ompT hsdS(rBmB) gal [malB$^{+}$] K-12(λS)* | Protein expression *in vivo*, protein purification | Merck, Darmstadt, Germany |
| C43(DE3) | F – *ompT hsdSB (rB- mB-) gal dcm* (DE3) | Protein expression *in vivo*, protein purification | [2] |

**Table S2: Plasmids used in this study**

Plasmids used for *in vivo* mRNA imaging lacked the Shine-Dalgarno sequence (SD).

| Name | Resistance/ Description | Application | Reference |
|---|---|---|---|
| pBad24 | Amp | Cloning vector | [3] |
| pET-19b | Amp | Cloning vector | Merck, Darmstadt, Germany |
| pRS1 | Amp | Cloning vector | [4] |
| pRS1-YohP-MS2.6x | Amp | In vivo and in vitro expression YohP | [5] |
| pRS1-YohP | Amp | In vivo and in vitro expression YohP without MS2 stem loop | This study |
| pBad24-YohP$_{His6}$, | Amp | In vivo expression YohP | [5] |
| pBad22-SecE(Y$_{GFP}$)G | Amp | In vivo expression | [6] |
| pBad22-SecE(YΔC4-C6$_{GFP}$)G | Amp | In vivo expression | This study |
| pBad22-SecE(YΔC4$_{GFP}$)G | Amp | In vivo expression | This study |
| pBad22-SecE(YΔC5$_{GFP}$)G | Amp | In vivo expression | This study |
| pBad22-SecE(YΔC6$_{GFP}$)G | Amp | In vivo expression | This study |
| pTRc99a-SecY$_{His}$EG | Amp | INV | [7] |
| pBad22-SecE(YΔC4-C6)G | Amp | INV | This study |
| pBad22-SecE(YΔC4)G | Amp | INV | This study |
| pBad22-SecE(YΔC5)G | Amp | INV | This study |
| pBad22-SecE(YΔC6)G | Amp | INV | This study |
| pTRC99a-YidC | Amp | INV | [8] |
| pTRC99a-FtsY | Amp | Protein purification | [9] |
| pTRC99a-Ffh | Amp | Protein purification | [10] |
| pT7/3a | Amp | In vitro transcription 4.5S RNA | [11] |
| pBad-MS2-venus | Amp | In vivo expression MS2-venus | This study |
| pSC.YohP-MS2.6x | Cm | In vivo expression of *yohP* mRNA | [5] |
| pSC.BglB-MS2.6x | Cm | In vivo expression of *bglB* mRNA | [5] |
| pSC.SecY-MS2.6x | Cm | In vivo expression of *secY* mRNA | This study |
| pSC.YohP-MS2-U-rich.6x | Cm | In vivo expression of U-rich *yohP* mRNA | This study |
| pSC.YohP-MS2-C-rich.6x | Cm | In vivo expression of C-rich *yohP* mRNA | This study |

| | | | |
|---|---|---|---|
| pSC.YohP-MS2-G-rich.6x | Cm | In vivo expression of G-rich *yohP* mRNA | This study |
| pSC.YohP-MS2-A-rich.6x | Cm | In vivo expression of A-rich *yohP* mRNA | This study |
| pSC.YohP(Δ4-30)-MS2.6x | Cm | In vivo expression of *yohP*(Δ4-30) mRNA | This study |
| pSC.YohP(Δ10-27)-MS2.6x | Cm | In vivo expression of *yohP*(Δ10-27) mRNA | This study |
| pSC.YohP(Δ46-60)-MS2.6x | Cm | In vivo expression of *yohP*(Δ46-60) mRNA | This study |
| pSC.YohP(4-30subst)-MS2.6x | Cm | In vivo expression of *yohP*(4-30subst) mRNA | This study |
| pSC.YohP(10-27subst)-MS2.6x | Cm | In vivo expression of *yohP*(10-27subst) mRNA | This study |
| pSC.YohP(46-60subst)-MS2.6x | Cm | In vivo expression of *yohP*(46-60subst) mRNA | This study |
| pSc.YohP.5’-3’ UTR ΔMS2.6X | Cm | In vivo expression of yohP mRNA without MS2 stem loop | This study |
| pBAD24.SecY-2TM_MS2-Venus | Amp | INV | This study |
| pRS1.YohP_MS2.6X | Amp | In vitro and in vivo study | This study |
| pRS1.YohP-MS2-U-rich.6x | Amp | In vitro and in vivo study | This study |
| pRS1.YohP-MS2-C-rich.6x | Amp | In vitro and in vivo study | This study |
| pRS1-YohP(ΔRBS).MS2.6X | Amp | In vitro protein synthesis | This study |
| pSc.YohP-MS2.6X-A-rich* | Cm | In vivo expression of *yohP* A-rich* mRNA | This study |
| pSc.YohP-MS2.6X-U-rich* | Cm | In vivo expression of *yohP* U-rich* mRNA | This study |
| pSc.YohP-MS2.6X-G-rich* | Cm | In vivo expression of *yohP* G-rich* mRNA | This study |
| pSc.YohP-MS2.6X-C-rich* | Cm | In vivo expression of *yohP* C-rich* mRNA | This study |
| pSc.YohP.5’-3’ UTR.MS2.6X | Cm | In vivo expression of *yohP* mRNA with 5’-3’-UTR | This study |

### Table S3: Oligonucleotides used in this study

| Name | Sequence 5’-3’ | Application |
|---|---|---|
| C4_trunc_fw | CATTTACCGCTGAAAGTGAATAGGC | Deletion of C4 loop SecY |
| C4_trunc_rev | GCGGCGTTGACCACGCTCAAC | Deletion of C4 loop SecY |
| C5_trunc_fw | AAAGTAATGACCCGCCTGACCCT | Deletion of C5 loop SecY |
| C5_trunc_rev | CTTTGCTGTTTCACGCGGGTTGAA | Deletion of C5 loop SecY |
| C6_trunc_fw | TTAGCGGCGGCGGGATCCGGA | Deletion of C6 loop SecY |
| C6_trunc_rev | GGACATCATCAGAGTTTGCACTTGA | Deletion of C6 loop SecY |
| pSc_Opening_Fw | GATGATATTTTAAGTAATTATAAC | Linearization of the plasmid pSc for Gibson assembly |
| pSc_Opening_rev | CTCAGGTCGACGTCCCATGGCCATTCG<br>AAT | Linearization of the plasmid pSc for Gibson assembly |
| U-rich_fw | AAT TCA GGA GGT CGA CCT GAG ATG<br>AAA ATT ATT CTT TGG GCT GTT TTG<br>ATT ATT TTC CTT ATT GGT CTTC TTG<br>TTG TTA CTG GTG TTT TTA AGA TGA<br>TTT TTT AAG | U-rich variant of YohP |
| U-rich_rev | AATTCAGGAGGTCGACCTGAGATGAA<br>AA TCATACTCTGGGCCGTACTGATCATC<br>TTC CTGATCGGGCTACTGGTGGTGA<br>CCGGCG TATTCAAGATGATATTCT<br>AAG | U-rich variant of YohP |
| C-rich_fw | AATTCAGGAGGTCGACCTGAGATGAAAAT<br>CATACTCTGGGCCGTACTGATCATCTTCCT<br>GATCGGGCTACTGGTGGTGACCGGCGTAT<br>TCAAGATGATATTCTAAG | C-rich variant of YohP |
| C-rich_rev | GATCCTTAGAATATCATCTTGAATACGCCGGT<br>CACCACCAGTAGCCCGATCAGGAAGATGATC<br>AGTACGGCCCAGAGTATGATTTTCATCTCAG<br>GTCGACCTCCTG | C-rich variant of YohP |
| G-rich_fw | GCC ATG GGA CGT CGA CCTG AGA TGA<br>AGA TTA TAC TGT GGG CGG T | G-rich variant of YohP |
| G-rich-rev | ATA ATT ACT TAA AAT ATC ATC TTA AAA<br>TAT CAT CTT AAA C | G-rich variant of YohP |
| A-rich_fw | CCA TGG GAC GTC GAC CTG AGA TGA<br>AAA TAA TAC TAT GGG CAG TAT TAA<br>TAA TAT TAG TAA CAG GAG TAT TTA<br>AAA TGA TAT TTG ATG ATA TTT TAA<br>GTA ATT A | A-rich variant of YohP |
| pRS1_Opening_Fw | TCGAGTAGCATAACCCCTTGGGGCCTCT<br>AAACGGG | Linearization of pRS1 for |

| | | |
|---|---|---|
| | | Gibson assembly |
| pRS1_Opening_rev | CATGGGGTATATCTCCTTCTTAAAGTTA AACAAAATTATTT | Linearization of pRS1 for Gibson assembly |
| YohP_MS2-6X_Fw | GAAGGAGATATACCCCATGATGAAAAT TATACTCTGGG | In vitro translation and pulse chase |
| YohP_MS2-6X_rev | GGGGTTATGCTACTCGAGCTAGAACTAT AGCTAGCATG | In vitro translation and pulse chase |
| U-rich_MS2.6X_fw | TATACCCCATGAATGAAAATTATTCTTT GGGC | In vitro translation and pulse chase |
| C-rich_MS2-6X | AGAAGGAGATATACCCCATGAATGAAA ATCATACTCTGGGCCGTAC | In vitro translation and pulse chase |
| YohP (Δ4-30)_fw | ATTTTCCTGATTGGGCTACTGGTGG | Imaging study |
| YohP (Δ4-30)_rev | CATCTCAGGTCGACGTCCCATG | Imaging study |
| YohP (Δ10-27)_fw | ATTATTTTCCTGATTGGGCTACTGGTGGT | Imaging study |
| YohP (Δ10-27)_rev | AATTTTCATCTCAGGTCGACGTCC CA | Imaging study |
| YohP (Δ46-60)_fw | GGCGTATTTAAGATGATATTTTAA GATGATATTTTAAGTAATTATAACCC | Imaging study |
| YohP (Δ46-60)_rev | CCCAATCAGGAAAATAATCAA TACAGCCCAGAGTATAATT | Imaging study |
| YohP(4-30 subst)_Fw | GGGCAGTTTTAATCATTTTCCTGATTGG GCTACTGGTGGTGACTGGCGT | Imaging study |
| YohP(4-30 subst)_rev | ACAGAATGATCTTCATCTCAGGTC GACGTCCCATGGCCA | Imaging study |
| YohP(10-27 subst)_rev | ATTTTCATCTCAGGTCGACGTCCC ATGG CCATTCGA | Imaging study |
| YohP(10-27 subst)_fw | TACACACTCGGATGCATCGATTATTTTC CTGATTGGGCTACTGGT | Imaging study |
| YohP(46-60 subst)_fw | AGTCACAGGCGTATTTAAGATGATATTT TAAGGCGTATTTAAG | Imaging study |
| YohP(46-60 subst)_rev | ACGAGAAGCCCAATCAGG AAAATAATC | Imaging study |
| pBAD24 Opening_fw | GGAGTCTGGTCGTCGTAAGATGGCTTCT AACTTTACTCAGTTCG | Linearization of pBAD24 for Gibson assembly |
| pBAD24 Opening_rev | CTAATCCCGGTTGTTTAGCCATGG GTACCATGGTGAATTCCTC | Linearization of pBAD24 for Gibson assembly |
| SecY_2TM_fw | CAGGAGGAATTCACCATGGTACCCATG GCTAAACAACCGGGATTAG | INV preparation |
| SecY_2TM_rev | CGA ACT GAG TAA AGT TAG AAG CCA TCT TAC GAC GAC CAG ACT CC | INV preparation |

| | | |
|---|---|---|
| A-rich*.6x_fw | CCA TGG GAC GTC GAC CTG AGA TGA AAA TTA TAC TCT GGG CTG TAT TAA TTA TTT TCC TAA TTG GGC TAC TGG<br>TAG TAA CAG GCG TAT TTA AAA TGA TAT TTT AAG TAA TTA TAA CCC GGG CCC T | Imaging study |
| U-rich*.6x_fw | CCA TGG GAC GTC GAC CTG AGA TGA AAA TTA TAC TTT GGG CTG TAT TGA TTA TTT TTC TTA TTG GTC TAC TGG TGG TGA CTG GTG TTT TTA AGA TGA TTT TTT AAG TAA TTA TAA CCC GGG CCC T | Imaging study |
| G-rich*.6x_fw | CCA TGG GAC GTC GAC CTG AGA TGA AAA TTA TAC TGT GGG CGG TGT TGA TTA TTT TCC TGA TTG GGC TAC TGG TGG TGA CTG GCG TGT TTA AGA TGA TAT TTT AAG TAA TTA TAA CCC GGG CCC | Imaging study |
| C-rich*.6x_fw | CCA TGG GAC GTC GAC CTG AGA TGA AAA TCA TAC TCT GGG CTG TAT TGA TCA TCT TCC TGA TCG GCC TAC TGG TGG TGA CTG GCG TCT TCA AGA TGA TCT TCT AAG TAA TTA TAA CCC GGG CCC T | Imaging study |
| YohP-5’-3’ UTR-MS2.6X_fw | GCC ATG GGA CGT CGA CCT GAG TAT ACA CTA AGT GAA TGA TAT CTT C | Imaging study |
| YohP-5’-3’ UTR-MS2.6X_rev | TAG GGC CCG GGT TAT AAT TAC AGC CCG GGG GTG CAG GGG GCG | Imaging study |
| FW-del MS2-6X | CGCGATCGATATCAGCGCTTTAAATTTGCGC | Imaging study |
| Re-del MS2-6x | GGTACCTTAGGATCCATATATAGGGCCC | Imaging study |

### Table S4: *yohP* probes used in this study

| Name | Sequence 5'-3' | Application |
|---|---|---|
| 1 | TCATTCACTTAGTGTATA | FISH |
| 2 | AAGATAAATCGGAAGATA | FISH |
| 3 | CGTTATCCATAAACGATT | FISH |
| 4 | AAAAACGAAGCCCTTTGC | FISH |
| 5 | TGCTGAATAAGTATAGGA | FISH |
| 6 | CGTTCCTTTATTTGTGAG | FISH |
| 7 | GAGTATAATTTTCATTGG | FISH |
| 8 | AATAATCAATACAGCCCA | FISH |
| 9 | CAGTAGCCCAATCAGGAA | FISH |
| 10 | AAATACGCCAGTCACCAC | FISH |
| 11 | ATTTTAAAATATCATCTT | FISH |
| 12 | ACCTGATGACATTAATTA | FISH |
| 13 | TATTCTCGTTATTTTCGG | FISH |
| 14 | GCAACAGGATGAGAGACT | FISH |
| 15 | AATGCACATGACAGGAGC | FISH |
| 16 | CCAGTGATTATATGAAGC | FISH |
| 17 | CCCTGCGCGCTCCTTGCG | FISH |
| 18 | CGGCGGCGATTGGCCGCC | FISH |
| 19 | GCCCGGGGGTGCAGGGGG | FISH |
| oligo probe for MS2 | CTGCAGACATGGGTGATCCTCATGT | Northern blot |

### References


1. Hanahan, D. (1983) Studies on transfromation of Escherichia coli with plasmids, *Journal of molecular biology*. **166**, 557-580.

2. Miroux, B. & Walker, J. E. (1996) Over-production of proteins in Escherichia coli: mutant hosts that allow synthesis of some membrane proteins and globular proteins at high levels, *Journal of molecular biology*. **260**, 289-98.

3. Guzman, L. M., Belin, D., Carson, M. J. & Beckwith, J. (1995) Tight regulation, modulation, and high-level expression by vectors containing the arabinose PBAD promoter, *Journal of bacteriology*. **177**, 4121-30.

4. Jauss, B., Petriman, N. A., Drepper, F., Franz, L., Sachelaru, I., Welte, T., Steinberg, R., Warscheid, B. & Koch, H. G. (2019) Non-competitive binding of PpiD and YidC to the SecYEG translocon expands the global view on the SecYEG interactome in E. coli, *The Journal of biological chemistry*.

5. Steinberg, R., Origi, A., Natriashvili, A., Sarmah, P., Licheva, M., Walker, P. M., Kraft, C., High, S., Luirink, J., Shi, W. Q., Helmstädter, M., Ulbrich, M. H. & Koch, H. G. (2020) Posttranslational insertion of small membrane proteins by the bacterial signal recognition particle, *PLoS biology*. **18**, e3000874.

6. Kuhn, P., Weiche, B., Sturm, L., Sommer, E., Drepper, F., Warscheid, B., Sourjik, V. & Koch, H. G. (2011) The bacterial SRP receptor, SecA and the ribosome use overlapping binding sites on the SecY translocon, *Traffic (Copenhagen, Denmark)*. **12**, 563-78.

7. Collinson, I., Breyton, C., Duong, F., Tziatzios, C., Schubert, D., Or, E., Rapoport, T. & Kuhlbrandt, W. (2001) Projection structure and oligomeric properties of a bacterial core protein translocase, *The EMBO journal*. **20**, 2462-71.

8. Welte, T., Kudva, R., Kuhn, P., Sturm, L., Braig, D., Muller, M., Warscheid, B., Drepper, F. & Koch, H. G. (2012) Promiscuous targeting of polytopic membrane proteins to SecYEG or YidC by the Escherichia coli signal recognition particle, *Molecular biology of the cell*. **23**, 464-79.

9. Braig, D., Bar, C., Thumfart, J. O. & Koch, H. G. (2009) Two cooperating helices constitute the lipid-binding domain of the bacterial SRP receptor, *Journal of molecular biology*. **390**, 401-13.

10. Braig, D., Mircheva, M., Sachelaru, I., van der Sluis, E. O., Sturm, L., Beckmann, R. & Koch, H. G. (2011) Signal sequence-independent SRP-SR complex formation at the membrane suggests an alternative targeting pathway within the SRP cycle, *Molecular biology of the cell.* **22**, 2309-23.

11. Wood, H., J, L. & Tollervey, D. (1992) Evolutionary conserved nucleotides within the E.coli 4.5S RNA are required for association with P48 in vitro and for optimal function in vivo, *Nucleic acids research.* **20**, 5919-5925.

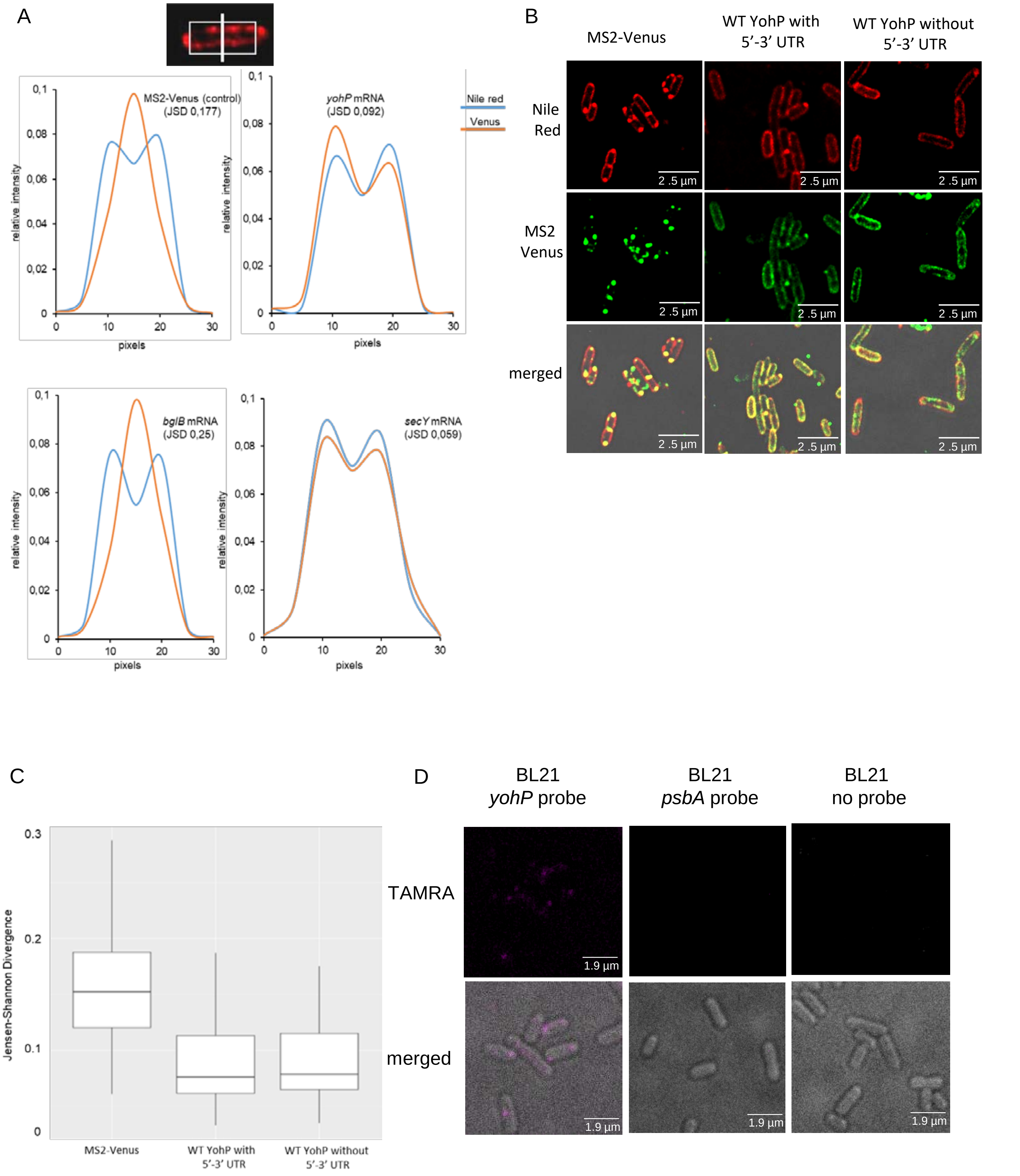


(Sarmah et al., Fig. S1)

A

```
WT-YohP          ATGAAAATTATACTCTGGGCTGTATTGATTATTTTCCTGATTGGGCTACTGGTGGTGACTGGCGTATTTAAGATGATATTTTAA
U-rich           ATGAAAATTATTCTTTGGGCTGTTTTGATTATTTTCCTTATTGGTCTTCTTGTTGTTACTGGTGTTTTTAAGATGATTTTTTAA
C-rich           ATGAAAATCATACTCTGGGCCGTACTGATCATCTTCCTGATCGGGCTACTGGTGGTGACCGGCGTATTCAAGATGATATTCTAA
A-rich           ATGAAAATAATACTATGGGCAGTATTAATAATATTCCTAATAGGACTACTAGTAGTAACAGGAGTATTTAAAATGATATTTTAA
G-rich           ATGAAGATTATACTGTGGGCGGTGTTGATTATTTTCCTGATTGGGCTGCTGGTGGTGACGGGGGTGTTTAAGATGATATTTTAA

U-rich*          ATGAAAATTATACTTTGGGCTGTATTGATTATTTTCTTATTGGTCTACTGGTGGTGACTGGTGTTTTTAAGATGATTTTTTAA
C-rich*          ATGAAAATCATACTCTGGGCTGTATTGATCATCTTCCTGATCGGCCTACTGGTGGTGACTGGCGTATTCAAGATGATCTTCTAA
A-rich*          ATGAAAATAATACTCTGGGCTGTATTAATTATTTTCCTAATTGGGCTACTGGTAGTAACAGGCGTATTTAAAATGATATTTTAA
G-rich*          ATGAAAATTATACTGTGGGCGGTGTTGATTATTTTCCTGATTGGGCTACTGGTGGTGACTGGCGTGTTTAAGATGATATTTTAA

yohP(Δ4-30)      ATG---------------------------ATTTTCCTGATTGGGCTACTGGTGGTGACTGGCGTATTTAAGATGATATTTTAA
yohP(Δ10-27)     ATGAAAATT------------------ATTATTTTCCTGATTGGGCTACTGGTGGTGACTGGCGTATTTAAGATGATATTTTAA
yohP(Δ46-60)     ATGAAAATTATACTCTGGGCTGTATTGATTATTTTCCTGATTGGG---------------GGCGTATTTAAGATGATATTTTAA

yohP(4-30subs)   ATGAAGATCATTCTGTGGGCAGTTTTAATCATTTTCCTGATTGGGCTACTGGTGGTGACTGGCGTATTTAAGATGATATTTTAA
yohP(10-27subs)  ATGAAAATTACACACTCGGATGCATCGATTATTTTCCTGATTGGGCTACTGGTGGTGACTGGCGTATTTAAGATGATATTTTAA
yohP(46-60subs)  ATGAAAATTATACTCTGGGCTGTATTGATTATTTTCCTGATTGGGCTTCTCGTAGTCACAGGCGTATTTAAGATGATATTTTAA
```

B

| Variant | **length** | **A** | **G** | **C** | **T** |
|---|---|---|---|---|---|
| WT | 84 | 21 | 20 | 9 | 34 |
| U-rich | 84 | 16 | 15 | 7 | 46 |
| C-rich | 84 | 21 | 20 | 18 | 25 |
| A-rich | 84 | 36 | 13 | 7 | 28 |
| G-rich | 84 | 17 | 28 | 7 | 32 |
| U-rich* | 84 | 19 | 18 | 6 | 41 |
| C-rich* | 84 | 20 | 19 | 17 | 28 |
| A-rich* | 84 | 28 | 15 | 9 | 32 |
| G-rich* | 84 | 19 | 24 | 8 | 33 |
| (Δ4-30) | 57 | 13 | 15 | 6 | 23 |
| (Δ10-27) | 66 | 18 | 15 | 6 | 27 |
| (Δ46-60) | 69 | 19 | 15 | 6 | 29 |
| (4-30 subs) | 84 | 20 | 21 | 10 | 33 |
| (10-27 subs) | 84 | 23 | 19 | 12 | 30 |
| (46-60 subs) | 84 | 22 | 17 | 11 | 34 |

C

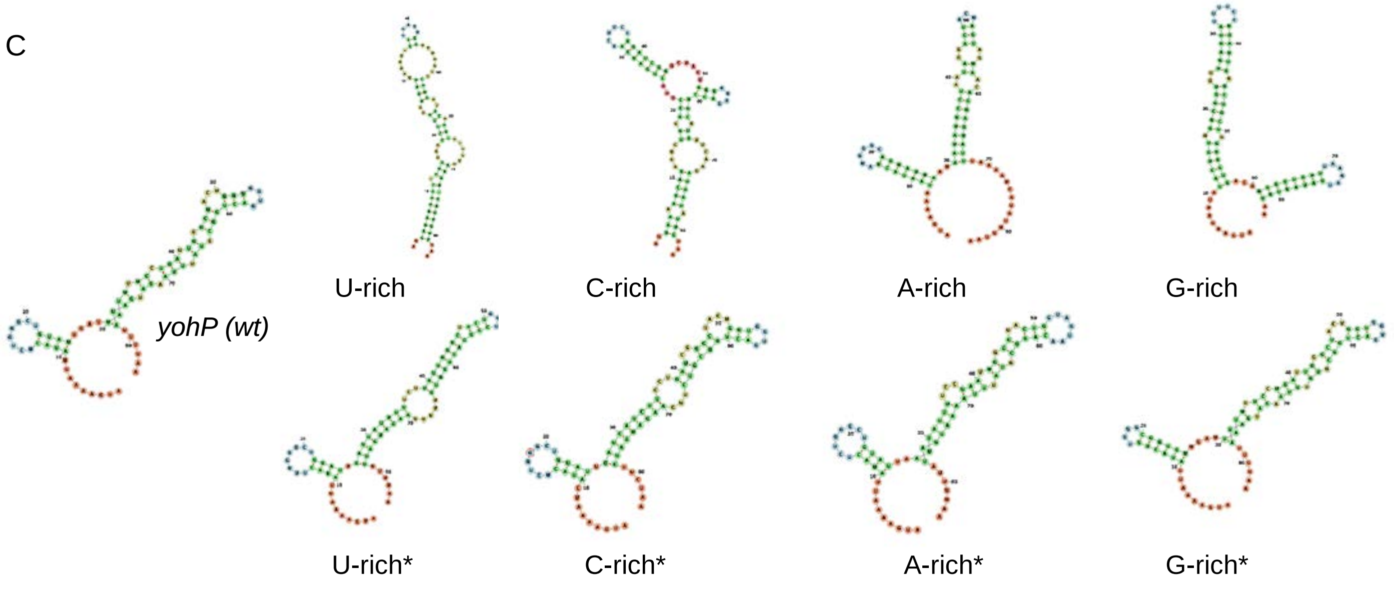


(Sarmah et al., Fig. S2)

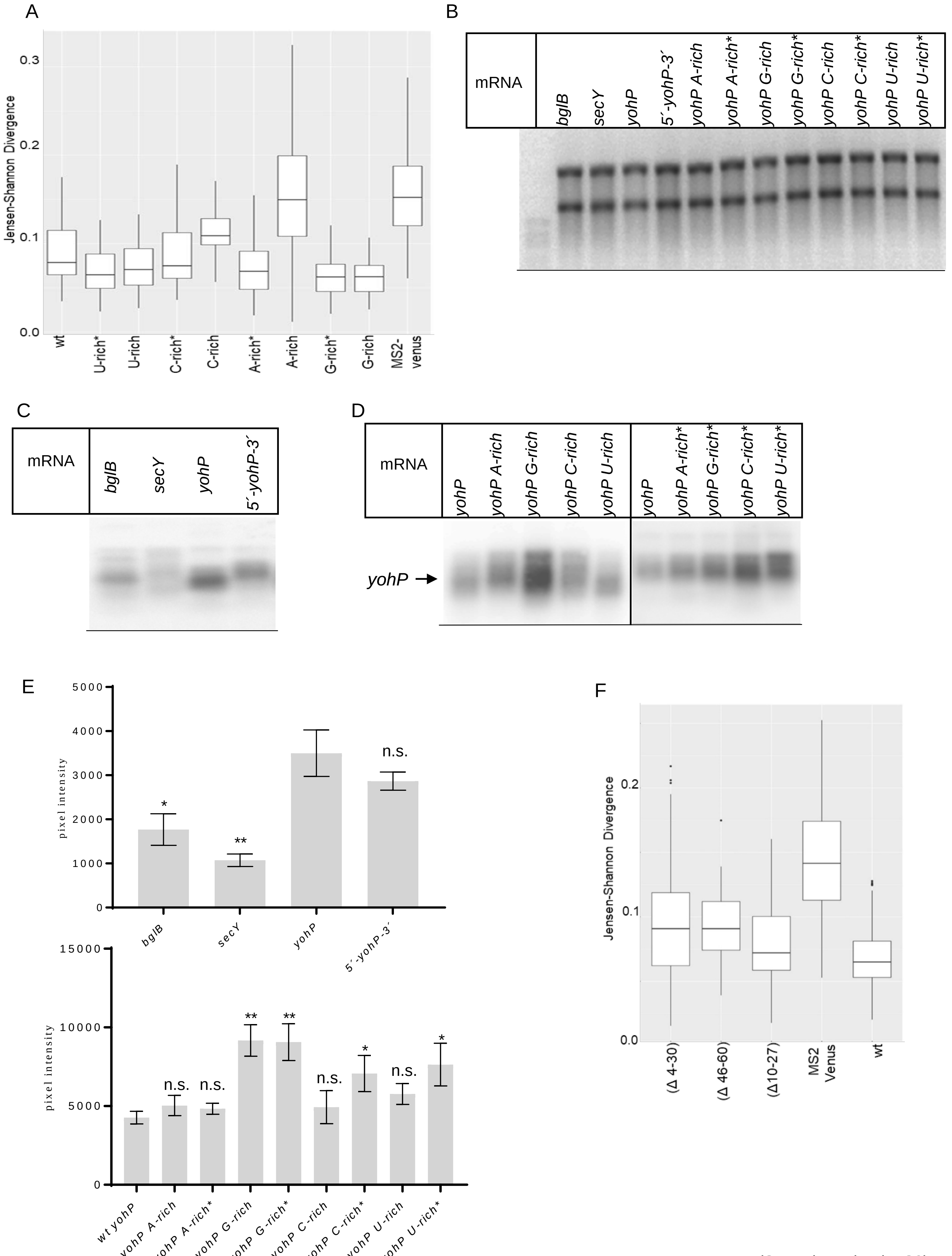


(Sarmah et al., Fig. S3)

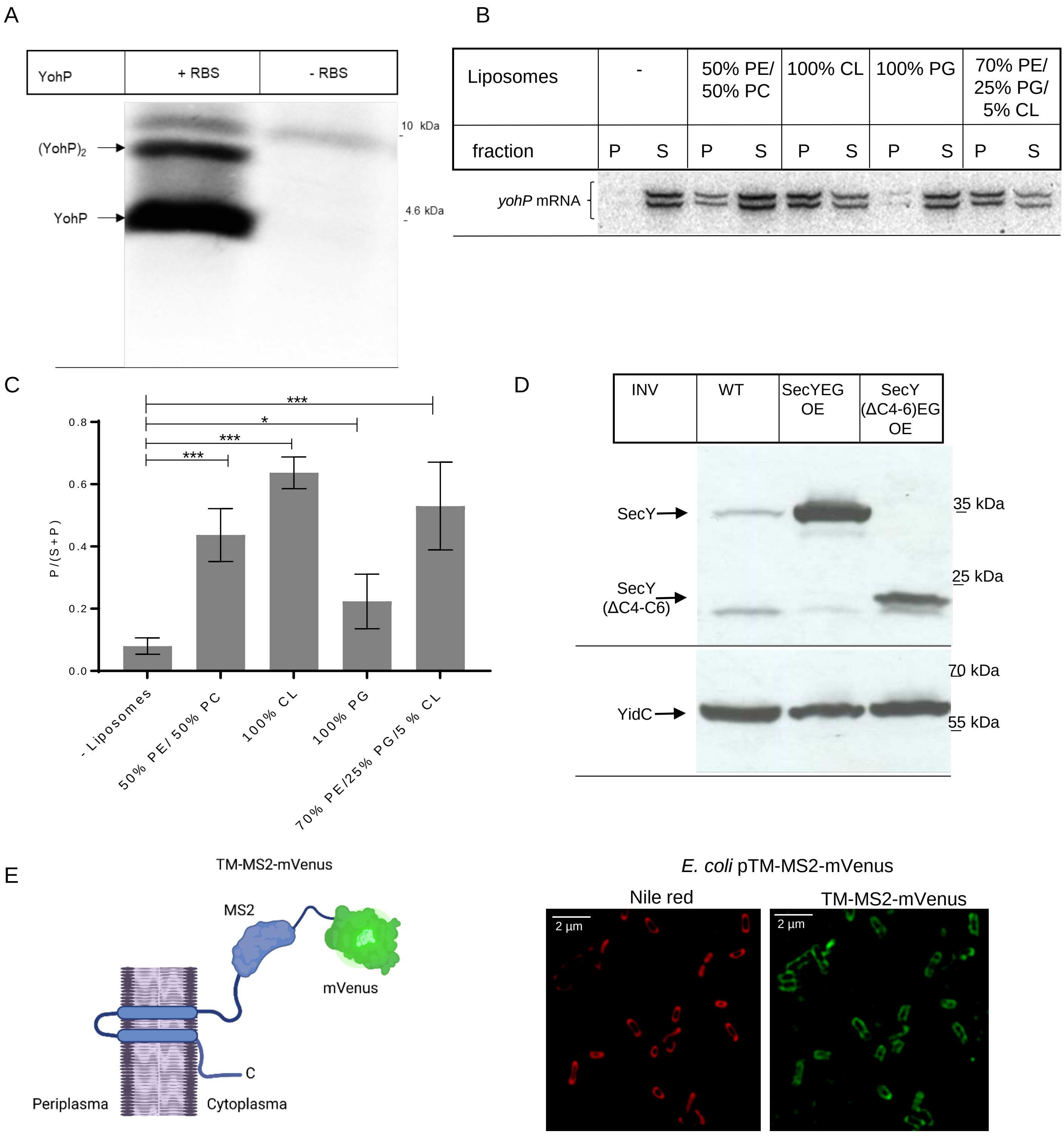


(Sarmah et al., Fig. S4)

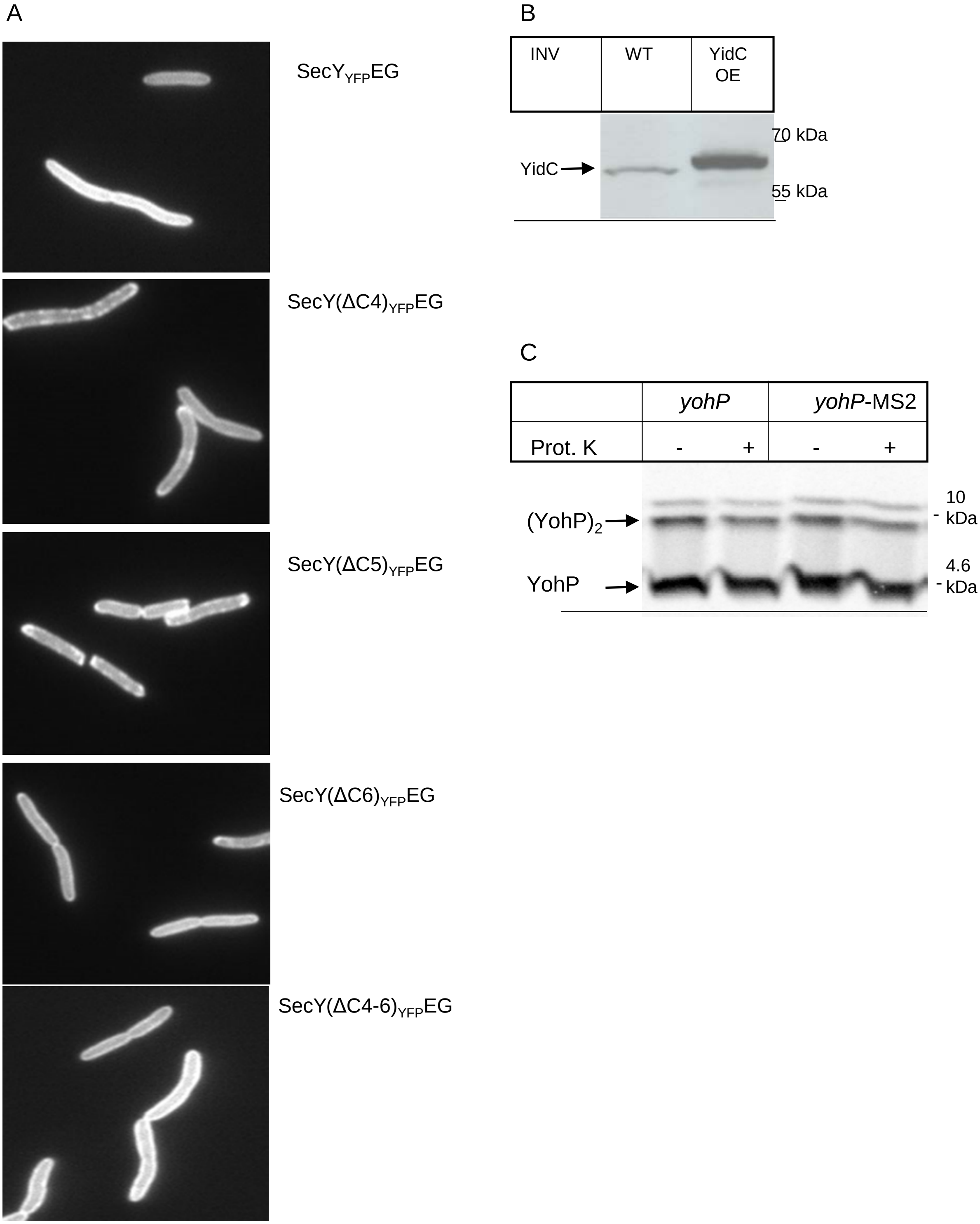


(Sarmah et al., Fig. S5 )

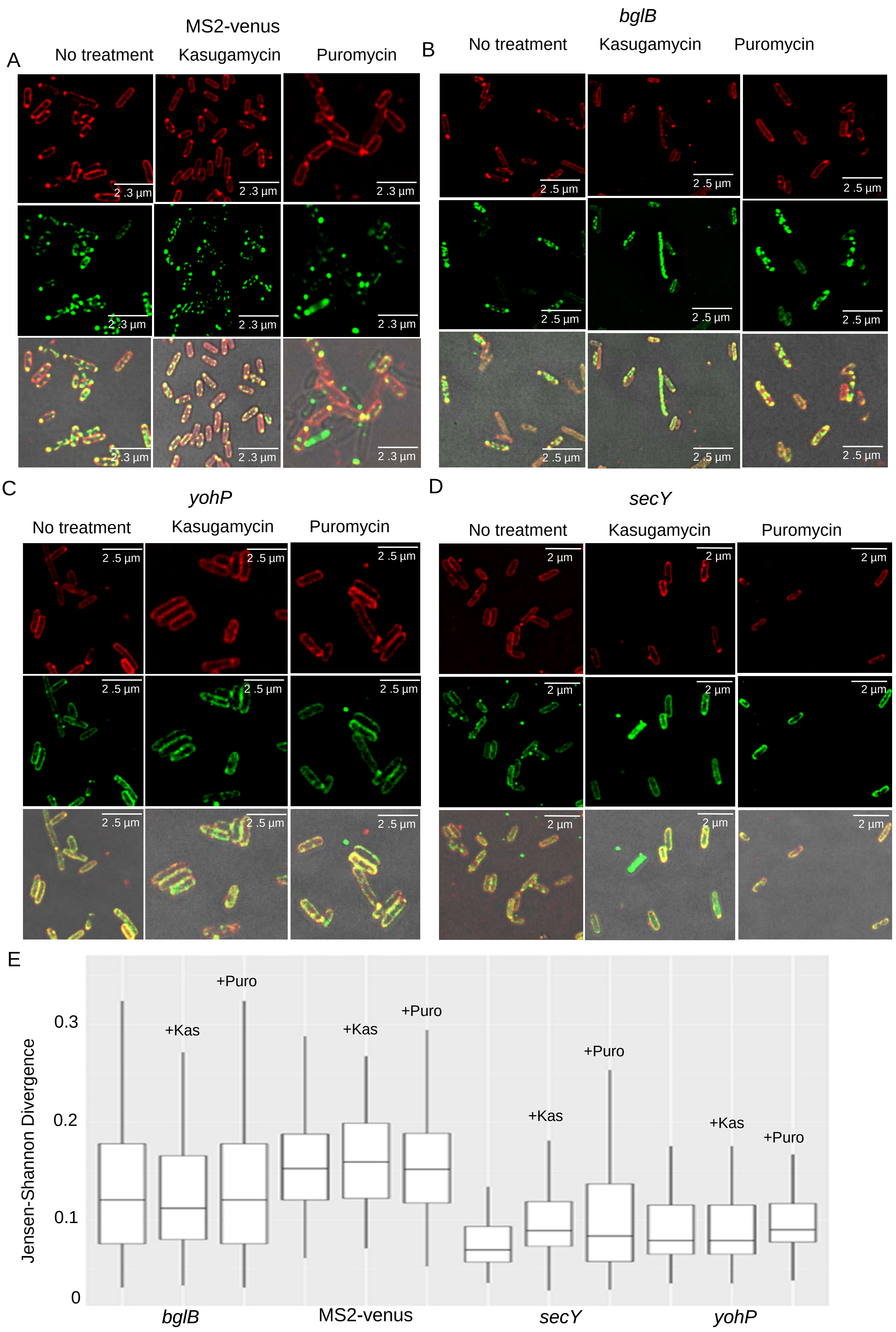


(Sarmah et al., Fig. S6 )

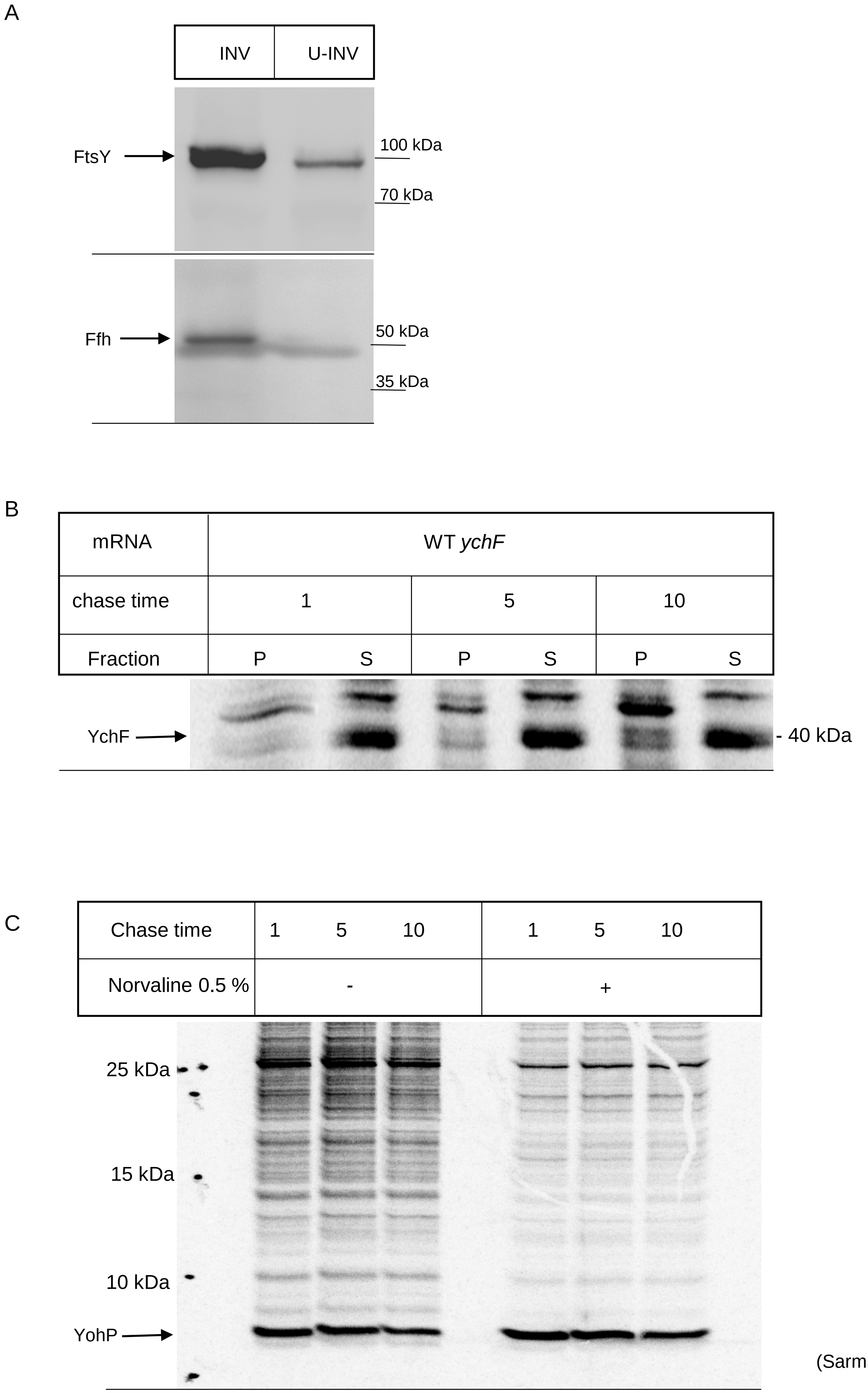


(Sarmah et al., Fig. S7 )
